# Architecture and dynamics of the jasmonic acid gene regulatory network

**DOI:** 10.1101/093682

**Authors:** Richard J. Hickman, Marcel C. Van Verk, Anja J.H. Van Dijken, Marciel Pereira Mendes, Irene A. Vos, Lotte Caarls, Merel Steenbergen, Ivo Van Der Nagel, Gert Jan Wesselink, Aleksey Jironkin, Adam Talbot, Johanna Rhodes, Michel de Vries, Robert C. Schuurink, Katherine Denby, Corné M.J. Pieterse, Saskia C.M. Van Wees

## Abstract

The phytohormone jasmonic acid (JA) is a critical regulator of plant growth and defense. To significantly advance our understanding of the architecture and dynamics of the JA gene regulatory network, we performed high-resolution RNA-Seq time series analyses of methyl JA-treated *Arabidopsis thaliana*. Computational analysis unraveled in detail the chronology of events that occur during the early and later phases of the JA response. Several transcription factors, including ERF16 and bHLH27, were uncovered as early components of the JA gene regulatory network with a role in pathogen and insect resistance. Moreover, analysis of subnetworks surrounding the JA-induced transcription factors ORA47, RAP2.6L, and ANAC055 provided novel insights into their regulatory role of defined JA network modules. Collectively, our work illuminates the complexity of the JA gene regulatory network, pinpoints to novel regulators, and provides a valuable resource for future studies on the function of JA signaling components in plant defense and development.

## INTRODUCTION

In nature, plants are subject to attack by a broad range of harmful pests and pathogens. To survive, plants have evolved a sophisticated immune signaling network that enables them to mount an effective defense response upon recognition of invaders. The phytohormone jasmonic acid (JA) and its derivatives are key regulators in this network and are typically synthesized in response to insect herbivory and infection by necrotrophic pathogens to activate specific defenses against these attackers^1^. Enhanced JA production mediates large-scale reprogramming of the plant’s transcriptome, which is influenced by the antagonistic or synergistic action of other hormones produced during parasitic interactions, such as salicylic acid (SA), ethylene (ET) or abscisic acid (ABA)^1–3^. The JA signaling network coordinates the production of a broad range of defense-related proteins and secondary metabolites, the composition of which is adapted to the environmental context and nature of the JA-inducing condition^1–3^.

In the past decade, major discoveries in the model plant *Arabidopsis thaliana* have greatly advanced our understanding of the JA signaling pathway. In the absence of an invader, activation of JA responsive gene expression is constrained by repressor proteins of the JASMONATE ZIM-domain (JAZ) family. Together with a co-repressor complex consisting of NOVEL INTERACTOR OF JAZ (NINJA) and TOPLESS, JAZs interact with and inhibit the action of transcription factors (TFs) that regulate JA-responsive genes. In response to pathogen or insect attack, the F-box protein CORONATINE INSENSITIVE1 (COI1) in the E3 ubiquitin-ligase Skip-Cullin-F-box complex SCF^CQI1^, which interacts with JAZs through their conserved C-terminal JA-associated (Jas) domain, perceives accumulation of bioactive JA-Isoleucine (JA-Ile). Upon perception of JA-Ile, JAZ repressor proteins are targeted for ubiquitination and subsequent proteasomal degradation^4–6^. This leads to the release of JAZ-bound TFs and subsequent induction of JA-responsive gene expression. Most JAZ pre-mRNAs are also subject to alternative splicing events that target the functionally important Jas motif. Alternative splicing events within the Jas domain can generate transcripts encoding truncated JAZ proteins that exhibit differential stability against JA-induced protein degradation. This increased functional diversity among JAZ proteins may increase regulatory complexity and achieve a more nuanced level of control over the JA pathway^7–9^.

Several groups of TFs are known to be important for regulation of the JA pathway. Upon degradation of JAZs, MYC2 acts in concert with the closely related bHLH TFs MYC3 and MYC4 in activating a large group of JA-responsive genes by directly targeting their promoters^10–12^. While current evidence indicates that MYC2, MYC3 and MYC4 act as master regulators of the onset of JA-responsive gene expression, additional factors are required for further fine-regulation of the dense JA-signaling circuitry. Several other bHLH TFs, such as JASMONATE-ASSOCIATED MYC2-LIKE1 (JAM 1)/bHLH017, JAM2/bHLH013, JAM3/bHLH003 and bHLH014 act redundantly to repress JA-inducible genes by competitive binding to *cis*-regulatory elements, possibly to control the timing and magnitude of the induced JA response^13–15^. Another important family of regulators that shape the JA response is the APETALA2/ETHYLENE RESPONSE FACTOR (AP2/ERF) family of TFs. AP2/ERF-type TFs, such as ERF1 and ORA59 (OCTADECANOID-RESPONSIVE ARABIDOPSIS AP2/ERF-domain protein 59), integrate the JA and ET response pathways and act antagonistically on MYC2,3,4-regulated JA-responsive genes^2, 16–18^. In general, AP2/ERF TF-regulated JA responses are associated with enhanced resistance to necrotrophic pathogens^16, 19^, whereas the MYC TF-regulated JA responses are associated with the wound response and defense against insect herbivores^20–22^.

Previously, several microarray-based transcriptome profiling studies revealed important information on the regulation of JA-responsive gene expression^23, 24^. However, because these studies analyzed this response at limited temporal resolution, much remains unknown about the architecture and dynamics of the JA gene regulatory network. Here, we performed an in-depth, high-throughput RNA sequencing (RNA-Seq) study in which we generated a high-resolution time series of the JA-mediated transcriptional response in developmental leaf number 6 of *Arabidopsis* plants. Computational analysis of the JA-induced transcriptional landscape provided insight into the architecture and dynamics of the JA gene regulatory network at an unprecedented level of detail. We accurately identified JA-induced expression profiles, and used these to predict and validate the function of several novel regulators of the JA immune regulatory network. We resolved the sequence of transcriptional events that take place following induction of the JA response, constructed a model of the JA gene regulatory network, and identified and validated subnetworks surrounding several JA-induced TFs. Furthermore, we uncovered differential *JAZ10* mRNA splice variant production as a component of the early responding JA transcriptional network.

## RESULTS

### A time course of MeJA-elicited transcriptional reprogramming

A key step towards a systems-level understanding of the architecture of the JA signaling network is to obtain comprehensive and accurate insight into the dynamic transcriptional reprogramming that takes place during the JA response. To go beyond earlier studies that analyzed the JA transcriptional response with a limited number of time points, we generated a high-resolution time series of JA-mediated transcriptional reprogramming in *Arabidopsis* leaves. Previously, dense time series experiments successfully deciphered gene regulatory networks underpinning a variety of biological processes^25–27^. Here, we profiled whole-genome transcriptome changes in *Arabidopsis* leaves treated with methyl JA (MeJA; that is readily converted to JA) or a mock solution over 15 consecutive time points within 16 h after treatment using RNA-Seq technology (Supplementary file 1A). Read counts were normalized for differences in sequencing depth between samples (Supplementary file 1B) and a generalized linear model was employed to identify genes whose transcript levels differed significantly over time between MeJA and mock treatments (see ref^28^ and methods for details). This yielded a set of 3611 differentially expressed genes (DEGs; Supplementary file 1C). Many of these DEGs were not previously described as MeJA responsive (Figure 1A), including 596 genes that are not represented on the ATH1 microarray used in earlier studies^23, 24^ A comparison of genes differentially expressed in our dataset revealed that 49% of these were also differentially expressed after feeding by the JA-inducing insect herbivore *Pieris rapae*^29^ (Figure 1A), indicating that our MeJA-induced time series captures a large proportion of biologically-induced transcriptional changes. Our high-resolution temporal transcriptome data captured a diverse set of expression profiles, including strongly transient expression changes, such as that of *JAR1* and *EDS1* (Figure 1B), which allows for modeling of biologically relevant JA-induced gene expression changes over time.

**Figure 1.**
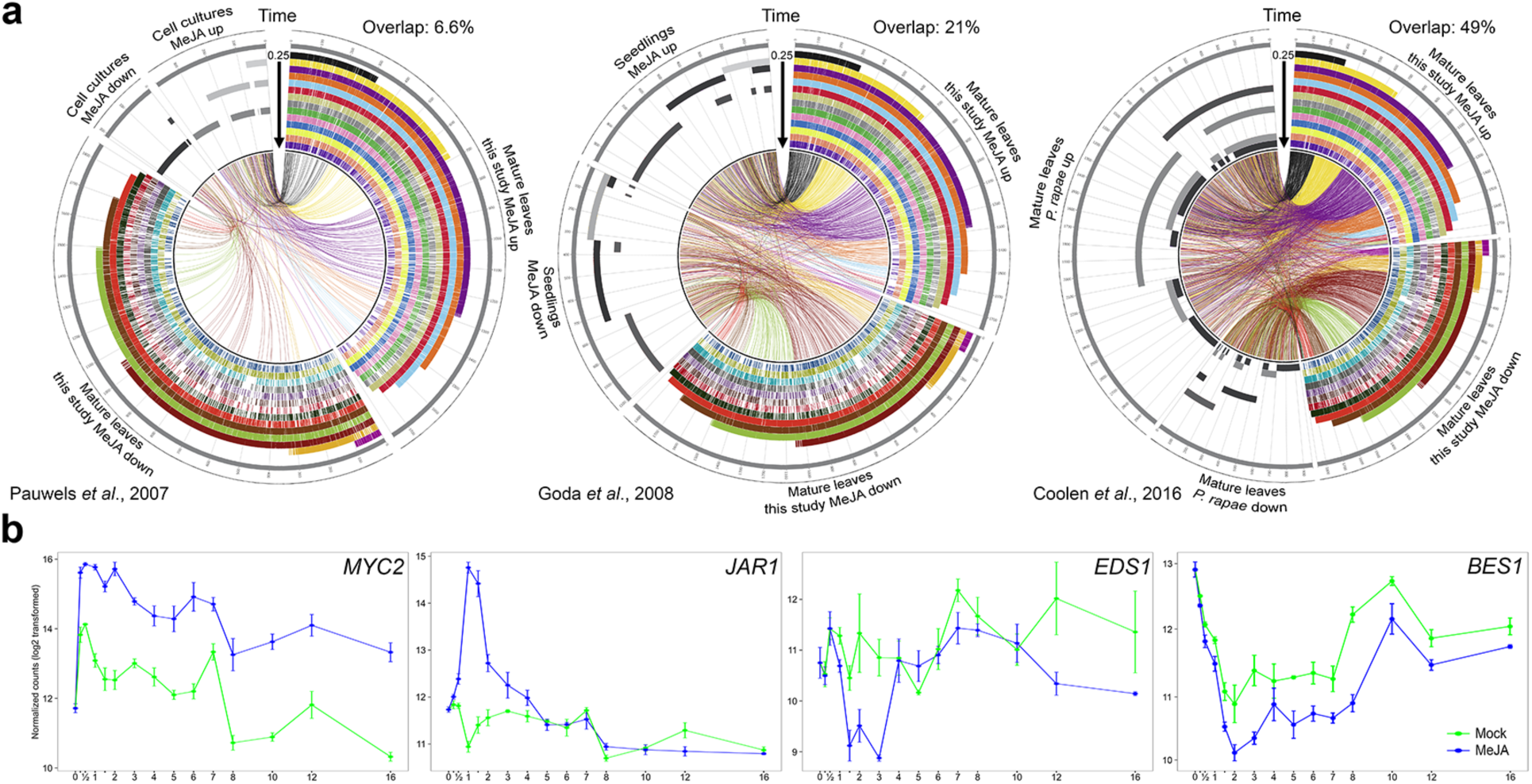
Temporal expression profiles following application of MeJA. **(a)** Circos plots of time series expression profiles from our MeJA time series experiment in comparison to previously published MeJA- or *P. rapae*-induced transcriptome data (Pauwels *et al*.^23^, Goda *et al*.^24^ and Coolen *et al*.^29^), as indicated underneath each plot). The stacked histograms indicate differential expression (colours indicate sampling time point from 0.25 h up to 16 h after treatment). Genes differentially expressed in both datasets are marked by connecting bands (colours indicate first time point of differential expression in our study). Each section within the circus plot represents a set of 100 DEGs. **(b)** Examples of expression profiles of selected JA pathway marker genes in our study. Y-axis, transcript abundance (log2-transformed read counts); x-axis, time (h) post application of MeJA; bars indicate SE.

### Process-specific gene clusters and novel defense regulators

To begin to decode the JA gene regulatory network, the time series-clustering algorithm SplineCluster was used to partition the set of 3611 DEGs into clusters of co-expressed genes. This yielded 27 distinct clusters (Figure 2A, Figure 2-figure supplement 1, Figure 2-source data 1), which broadly fall into two major groups: those that show increased expression in response to application of MeJA (cluster 1-14), and those that exhibit reduced expression (cluster 15-27). To facilitate the use of the expression data for the *Arabidopsis* community, a searchable (gene ID) figure has been made available that visualizes co-expression relationships in time for all DEGs in the individual clusters (Supplementary file 1D). The genes in each cluster were tested for overrepresented functional categories using Gene Ontology (GO) term enrichment analysis to investigate the biological significance of the distinct dynamic expression patterns (Figure 2-source data 2). This analysis showed that clusters representing up-regulated genes are, as expected, overrepresented for functional terms associated with JA defense responses. Broad annotations such as ‘Response to wounding’ and ‘Response to herbivory’ are present in multiple up-regulated clusters, while in contrast the more specific functional categories are linked specifically to single clusters. For example, cluster 6 is specifically overrepresented for the annotation term ‘Anthocyanin-containing compound biosynthetic process’, cluster 8 for ‘Tryptophan biosynthetic process’, and cluster 14 for ‘Glucosinolate biosynthetic process’. Each of these clusters contains many of the genes previously implicated in these secondary metabolite biosynthesis pathways, but also uncharacterized genes which may have an important function in these specific processes (Figure 2-source data 2). This indicates that the dynamic expression profiles generated in this study possess information that is sufficiently detailed to identify process-specific sectors in the JA gene regulatory network that are subject to distinct regulation.

**Figure 2.**
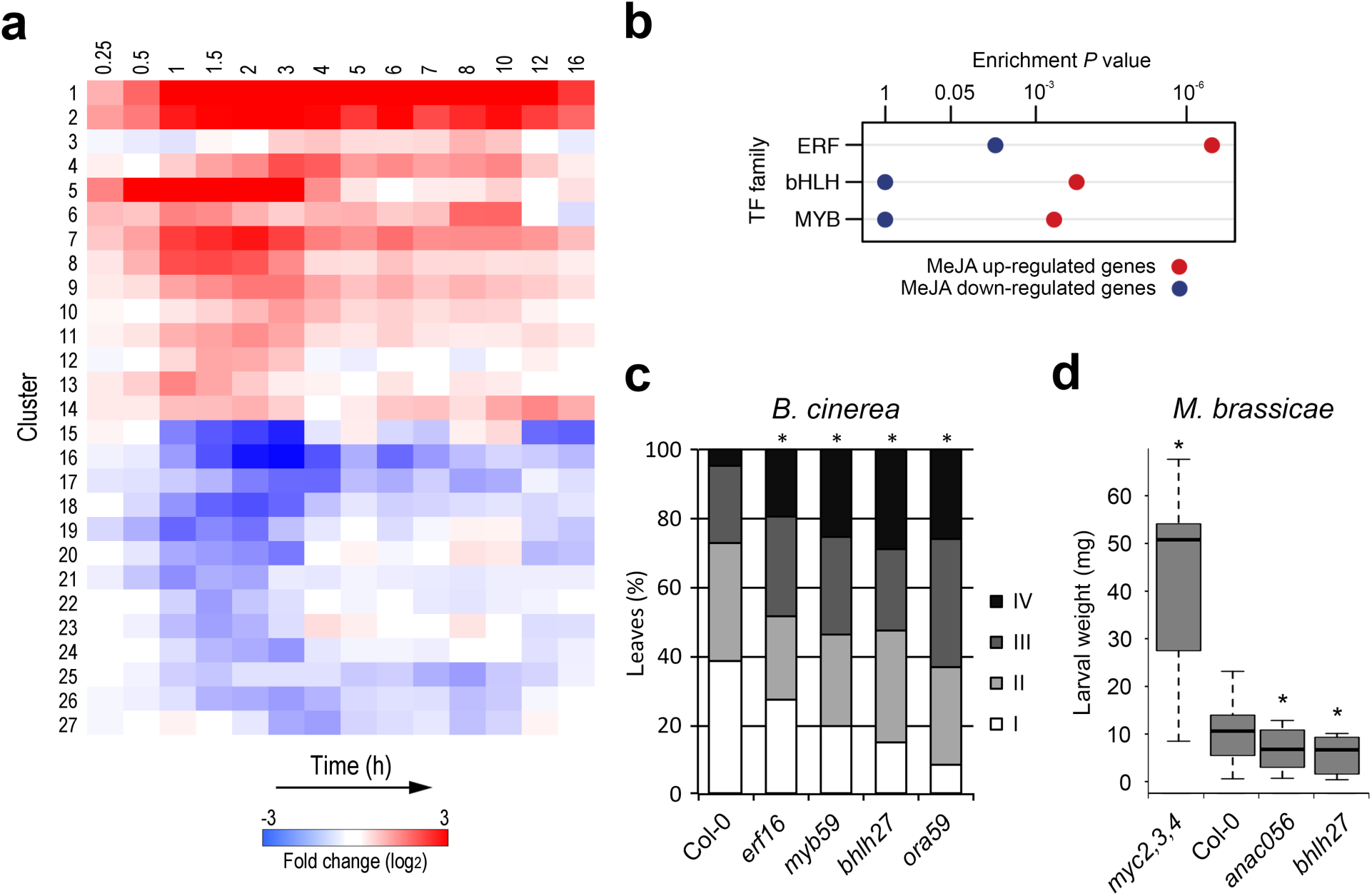
Clustering of co-expressed genes in the JA gene regulatory network and identification of novel components of JA-dependent resistance. **(a)** The set of 3611 genes showing differential expression in *Arabidopsis* leaves following exogenous application of MeJA were partitioned into 27 distinct co-expressed gene clusters using SplineCluster. The heatmap shows the mean gene expression profile for each cluster, with red and blue indicating up-regulation and down-regulation of expression (log2-fold change (MeJA/mock)), respectively. **(b)** Significantly overrepresented TF families within clusters of genes up-regulated (clusters 1-14; red) or down-regulated (clusters 15-27; blue) in response to MeJA treatment (hypergeometric test; *P* ≤ 0.001). **(c)** Quantification of disease symptoms of wild-type Col-0, highly susceptible ERF TF mutant *ora59*, and T-DNA insertion lines for selected genes *ERF16, MYB59*, and *bHLH27* at 3 days after inoculation with B. cinerea. Disease severity of inoculated leaves was scored in four classes ranging from restricted lesion (class I), non-spreading lesion (class II), spreading lesion (class III), up to severely spreading lesion (class IV). The percentage of leaves in each class was calculated per plant (n=20). Asterisk indicates statistically significant difference from Col-0 (Chi-squared test; *P* ≤ 0.05). **(d)** Performance of *M. brassicae* larvae on Col-0, highly susceptible triple bHLH TF mutant *myc2,3,4* and T-DNA insertion lines for selected genes *ANAC056* and *bHLH27*. The larval fresh weight was determined after 8 days of feeding. Asterisk indicates statistically significant difference from Col-0 (two-tailed Student’s *t* test for pairwise comparisons; *P* ≤ 0.05; n=30; error bars are SE).

Since TFs are the main drivers of transcriptional networks, we mapped the TF families that are enriched in the 27 clusters of MeJA-responsive DEGs. Within the up-regulated clusters, genes encoding members of the bHLH, ERF and MYB TF families were most significantly overrepresented (Figure 2B), suggesting that these TF families dominate the onset of JA-induced gene expression. The early up-regulated gene clusters 1 and 2 (61 and 165 genes, respectively) contained an enrichment for known JA-related genes such as the herbivory markers *VSP1* and *VSP2,* as well as the regulators *JAZ1, 2, 5, 7, 8, 9, 10* and *13, MYC2, ANAC019, ANAC055, RGL3,* and *JAM1*^30^ In addition, TF genes with no previously reported roles in the JA response pathway are present in these clusters, which implies that they may also have regulatory functions in the JA response. To test this, we selected 7 uncharacterized TF genes from clusters 1 and 2 and supplemented this set with 5 uncharacterized TF genes from other clusters, displaying a similarly rapid response to MeJA treatment. The respective *Arabidopsis* T-DNA knockout lines were functionally analyzed for their resistance against the necrotrophic fungus *Botrytis cinerea* and the generalist insect herbivore *Mamestra brassicae*, which are both controlled by JA-inducible defenses^2^. Mutants in the TF genes *bHLH27, ERF16* and *MYB59* displayed a significant increase in disease susceptibility to *B. cinerea* compared to wild-type *Arabidopsis* Col-0, approaching the disease severity levels of the highly susceptible control mutant *ora59* (Figure 2C; full results in Figure 2-figure supplement 2 and 4). Weight gain of *M. brassicae* larvae was significantly reduced on mutants of *ANAC056* and *bHLH27,* while on none of the tested mutants larval weight was enhanced, as was the case on the susceptible control mutant *myc2,3,4* (Figure 2D; full results in Figure 2-figure supplement 3 and 4). Thus, for 4 of the 12 tested MeJA-responsive TF genes a predicted role in the JA response could be functionally validated for either *B. cinerea* or *M. brassicae* resistance, demonstrating the utility of high-resolution time series to uncover co-expressed genes with novel functions in the JA pathway.

### Enrichment of TF DNA-binding motifs

TFs regulate gene expression by binding to *cis*-regulatory elements of target genes in a sequence specific manner. Mapping of regulatory DNA motifs that are associated with dynamic MeJA-responsive gene expression profiles can aid in the reconstruction of JA gene regulatory networks. Therefore, we investigated which *Arabidopsis* TF-binding site motifs are overrepresented within the promoters of the co-expressed MeJA-responsive DEGs. To this end, we utilized the recently characterized DNA-binding specificities for 580 *Arabidopsis* TFs from studies using protein-binding microarrays (PBMs), which identified 399 DNA motifs^31, 32^. Firstly, we screened for overrepresentation of these motifs in the unions of up- and down-regulated gene clusters, respectively (Figure 3A). Motifs corresponding to DNA-binding sites of bHLH, bZIP, ERF and MYB TFs are clearly overrepresented in the group of up-regulated genes, while WRKY and TCP TF specific motifs are markedly overrepresented in the down-regulated genes. Members of the WRKY TF family and their cognate *cis*-elements are key regulators of the SA response pathway^33^, suggesting that WRKYs are important targets in the transcriptional repression of the SA pathway by MeJA treatment. Secondly, we analyzed motif enrichment within each of the 27 clusters of co-expressed genes (Figure 3B). To increase the chance of discovering nuanced sequence motifs among the genes in these clusters, we supplemented the known motif analysis (Figure 3-source data 1) with *de novo* motif discovery (Figure 3-source data 2, 3). This revealed promoter elements that are selectively enriched in specific clusters, offering a more precise link between motifs and cluster-specific gene expression patterns. Strikingly, while motifs that correspond to bHLH-binding sites are enriched in the majority of the up-regulated gene clusters, ERF- and MYB-binding motifs are only overrepresented in a small selection of the up-regulated clusters that are associated with specific biological processes (e.g. clusters 6, 9, 13 and 14; Figure 3-source data 1). For example, clusters 6 and 14, which are enriched for GO terms describing distinct secondary metabolite biosynthesis pathways, are enriched for different *(de novo)* predicted MYB DNA-binding motifs (Figure 3B). These findings suggest that bHLH TFs and their DNA-binding sites are essential components in activation of the majority of the MeJA-inducible genes, while ERF and MYB TFs have more specialized roles in modulating the expression of dedicated sets of target genes.

**Figure 3.**
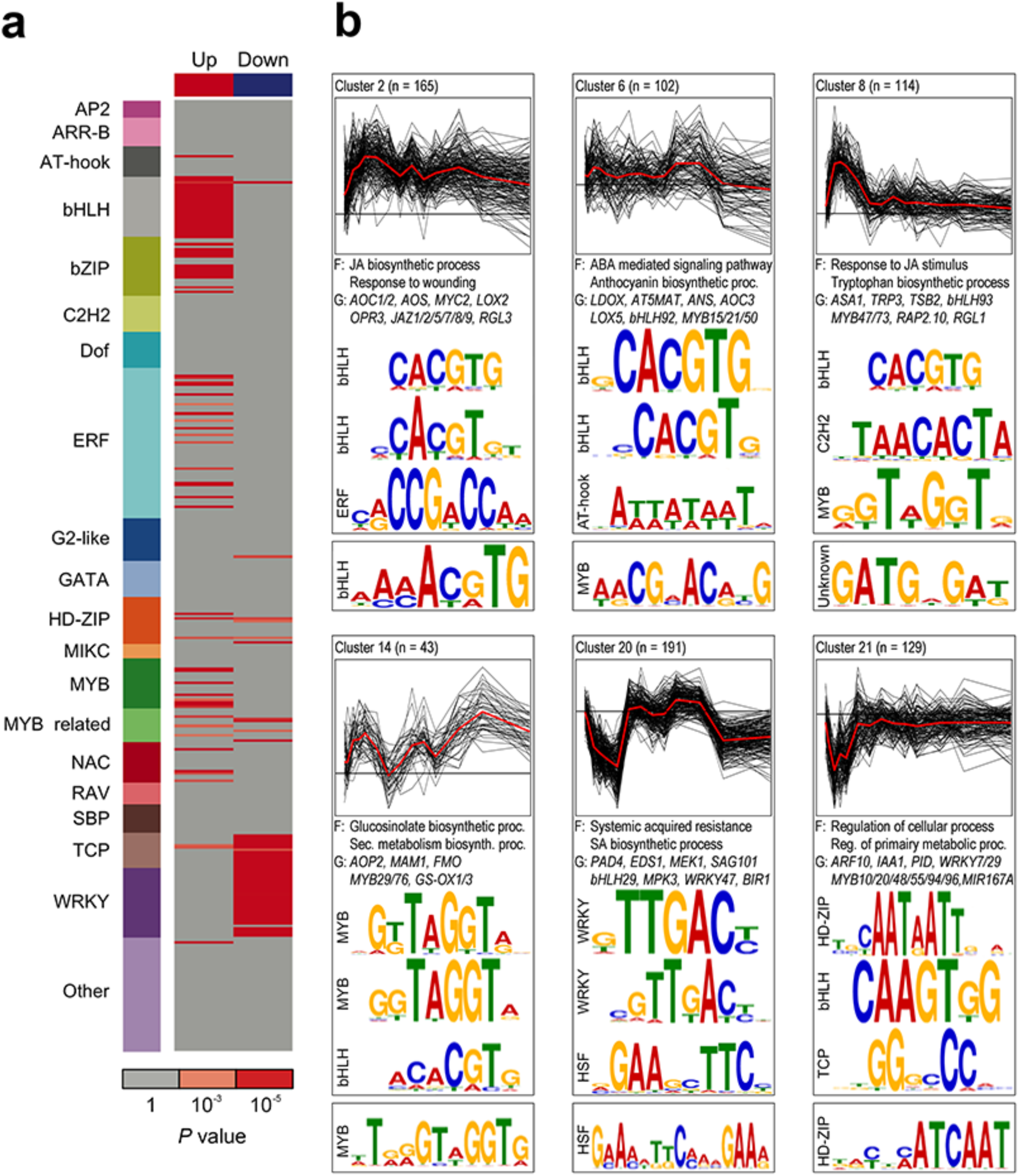
Enriched *cis*-regulatory motifs and functional categories in MeJA-responsive gene co-expression clusters. **(a)** Overrepresentation of 399 known TF DNA-binding motifs within the unions of up-regulated and downregulated genes. Rows indicate motifs and are colored by corresponding TF family. Red boxes indicate a motif that is significantly overrepresented (cumulative hypergeometric distribution). **(b)** Representative co-expression clusters with overrepresented TF DNA-binding motifs. Top: Profiles of log2-fold change in gene expression (MeJA/mock), with mean profile (red) and cluster size (n). Selected overrepresented functional categories (F) and representative genes (G) are denoted. Sequence logo depiction of selected known (middle) and de novo-derived (bottom) motifs that are significantly overrepresented.

### Chronology of MeJA-mediated transcriptional reprogramming

Next, we utilized the temporal information in our RNA-Seq time series to resolve the chronology of gene expression events in the JA gene regulatory network. First, we divided the genes in sets of up- and down-regulated DEGs and sorted them according to the time at which they first become differentially expressed (Figure 4-figure supplement 1; see methods for details). From this analysis, it became clear that a massive onset of gene activation precedes that of gene down-regulation, and that different waves of coordinated gene expression changes can be identified in the time series. The majority of all DEGs become first differentially expressed within 4 h after MeJA treatment, which supports the hypothesis that shallow network architectures favor rapid responses to external signals^34^. Up- and down-regulated DEGs were then further separated into two additional sets based on their predicted function as transcriptional regulators (termed regulator genes) or as having a different function (termed regulated genes; Supplementary file 1C). We were specifically interested in identifying time points where coordinated switches in transcriptional activity take place. We reasoned that pairs of adjacent time points that display a weaker correlation indicate important points of coordinated switches in transcriptional activity (see methods for details; Figure 4-figure supplement 2). Therefore, within each of the four mutually exclusive gene sets, we examined the pairwise correlations of expression levels between all pairs of time points. Clustering of the resulting correlation matrices revealed six distinct phases in the transcriptional activation, and four phases in the transcriptional repression (Figure 4A). The first two phases of up-regulation (Phase Up1 and Up2) start within 0.5 h after MeJA treatment in the set of regulator genes, while at 1.5 h a third phase of up-regulation of regulator genes ensues (phase Up4). For the regulated genes the first phase of up-regulation starts at 1 h after MeJA treatment (phase Up3), which is clearly later than the first onset of the regulator genes. A similar sequence of events can be observed in the down-regulated regulator and regulated genes, although the start is delayed compared to the activation of up-regulated genes.

**Figure 4.**
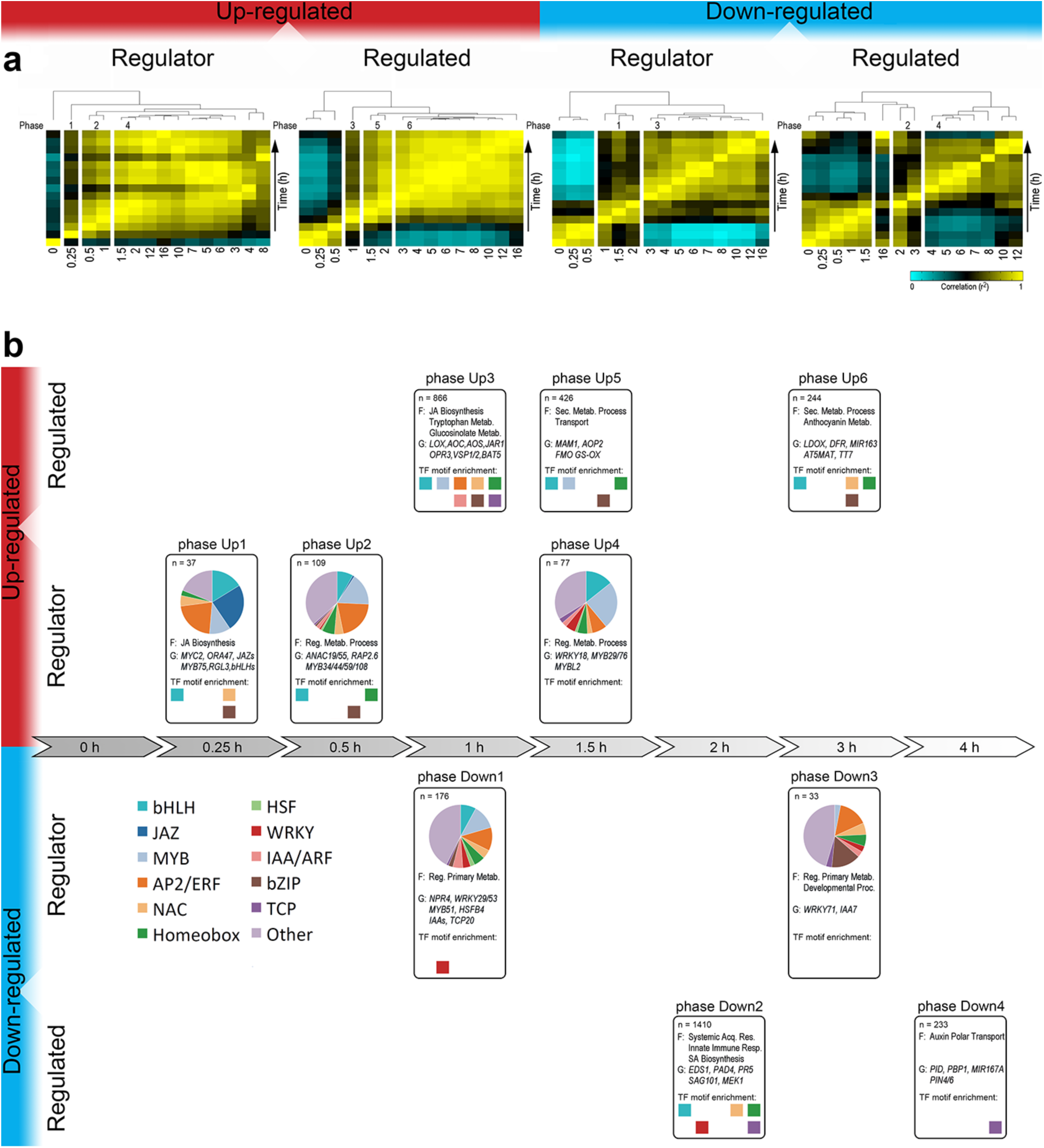
Chronology of changes in the MeJA-triggered gene regulatory network. **(a)** Phasing of MeJA-induced transcriptional changes. DEGs were divided into four sets according to their function as regulator or non-regulator (regulated), and their expression pattern being up- (red) or down-regulated (blue) over time. For each set of genes, a correlation matrix of gene transcription counts between all pairs of time points was computed using Pearson’s correlation metric. Shown are the dendrograms produced by hierarchical clustering of the transcriptome correlation matrices (yellow, high correlation; cyan, low correlation). Time is in hours. **(b)** Analysis of the major transcriptional phases in the JA gene regulatory network. Transcriptional phases are indicated by boxes, aligned on the timeline. DEGs are assigned to the phases according to the time point where they become first differentially expressed; indicated are overrepresented functional categories (F) and representative genes (G). Colored squares indicate known TF DNA-binding motifs overrepresented in gene promoters (hypergeometric distribution; *P* ≤ 0.001). Pie charts indicate the proportion of TF gene families.

To explore the biological significance and directionality in the regulation of the identified transcriptional phases in the JA gene regulatory network, all DEGs were assigned to the phase in which they first became differentially expressed (see methods for details; Figure 4-figure supplement 2). The resulting gene lists of the 10 transcriptional phases were tested for overrepresentation of functional categories and promoter motifs (Figure 4B; Figure 4-source data 1-4). Phase Up1 represents the immediate transcriptional response with genes encoding bHLHs, JAZs, MYBs and ERFs, and other genes associated with JA biosynthesis. These early regulator genes may play a role in the induction of other regulator genes present in phases Up2 and 4, and of regulated genes present in phases Up3, 5 and 6, which are linked to defense responses such as glucosinolate, tryptophan and anthocyanin biosynthesis (Figure 4B; Figure 4-source data 2). In support of this, in the DEGs of phase Up3, DNA motifs that can be bound by TFs transcribed in previous phases Up1 and 2, like bHLH-, ERF- and MYB-binding motifs, are enriched. In phase Up3, genes involved in JA biosynthesis are also enriched. Overall, induction of the JA pathway shows a clear chronology of up-regulated gene expression events, starting with the activation of genes encoding specific classes of TFs and of JA biosynthesis enzymes, followed by genes encoding enzymes involved in the production of defensive secondary metabolites.

The first wave of transcriptional repression by MeJA is also marked by genes encoding transcriptional regulators, and begins at 1 h after MeJA treatment with phase Down1, after which phases Down2, 3 and 4 follow at 2, 3 and 4 h after MeJA treatment, respectively (Figure 4B; Figure 4-source data 1). These groups of down-regulated genes highlight the antagonistic effects of JA on other signaling pathways. Phase Down1 for instance, is characterized by the repression of different defense-related genes such as *NPR4* and *MYB51,* which encode regulators that promote SA responses and indolic glucosinolate biosynthesis, respectively^35, 36^. Accordingly, *MYC2,* which is induced by MeJA in our dataset, was previously shown to suppress the accumulation of indolic glucosinolates^10^. Phase Down2 is also enriched for genes associated with SA-controlled immunity, including the key immune-regulators *EDS1* and *PAD4*^3^. In line with these observations, the promoters of the MeJA-repressed genes are overrepresented for DNA-binding motifs for WRKY TFs, suggesting that WRKY-regulated gene expression may be suppressed by MeJA treatment, which correlates with the antagonism of JA on SA signaling. Later phases of transcriptional repression (phases Down3 and 4) are marked by an overrepresentation of genes related to growth and development, including auxin signaling, and an enrichment of DNA motifs recognized by TCP TFs, which conceivably reflects an effort by the plant to switch energy resources from growth to defense^38^. A general observation that can be made from this chronological analysis of the JA gene regulatory network is that activation/repression is controlled by relatively short transcriptional cascades yielding distinctive transcriptional signatures.

### Inference of regulatory interactions reveals key regulators of local JA subnetworks

Next, we made use of the TF DNA-binding motif information of the genes in the temporally separated transcriptional phases to construct a gene regulatory network that predicts directional interactions between the JA responsive TF genes and all genes associated with the different transcriptional phases (Figure 5-source data 1). The JA gene regulatory network generated via this analysis is shown in Figure 5A, in which a differentially expressed TF gene (represented by a circular node in the network) is connected by an edge to a transcriptional phase (represented by a square in the network) when the corresponding DNA-binding motif is overrepresented in that phase. The network shows that the TFs in the network are predicted to regulate expression of genes at either single or multiple transcriptional phases. We were particularly interested in using the network to predict regulatory interactions between TFs assigned to the phase representing the first wave of transcriptional activity (Phase Up1), as TFs that become active in this phase are likely key regulators of subsequent transcriptional activity. Phase Up1 contains the TFs MYC2 and JAM1, which are among the most active TFs as their cognate DNA-binding motifs (both share the same consensus, CACGTG) are enriched in the promoters of genes assigned to a large fraction of the up-regulated transcriptional phases. This prediction is in line with recent reports suggesting that the positive regulator MYC2 and the negative regulator JAM1 cooperate to balance JA responses by competitive binding to their shared target sequences^13, 14^. What determines the different timing in transcriptional activation by these TFs awaits further investigation. Phases Up1 and Up2 contain the TF genes *bHLH27, ERF16, ANAC056* and *MYB59,* of which corresponding mutants show altered resistance levels to *B. cinerea* infection and *M. brassicae* infestation (Figure 2C, 2D).

**Figure 5.**
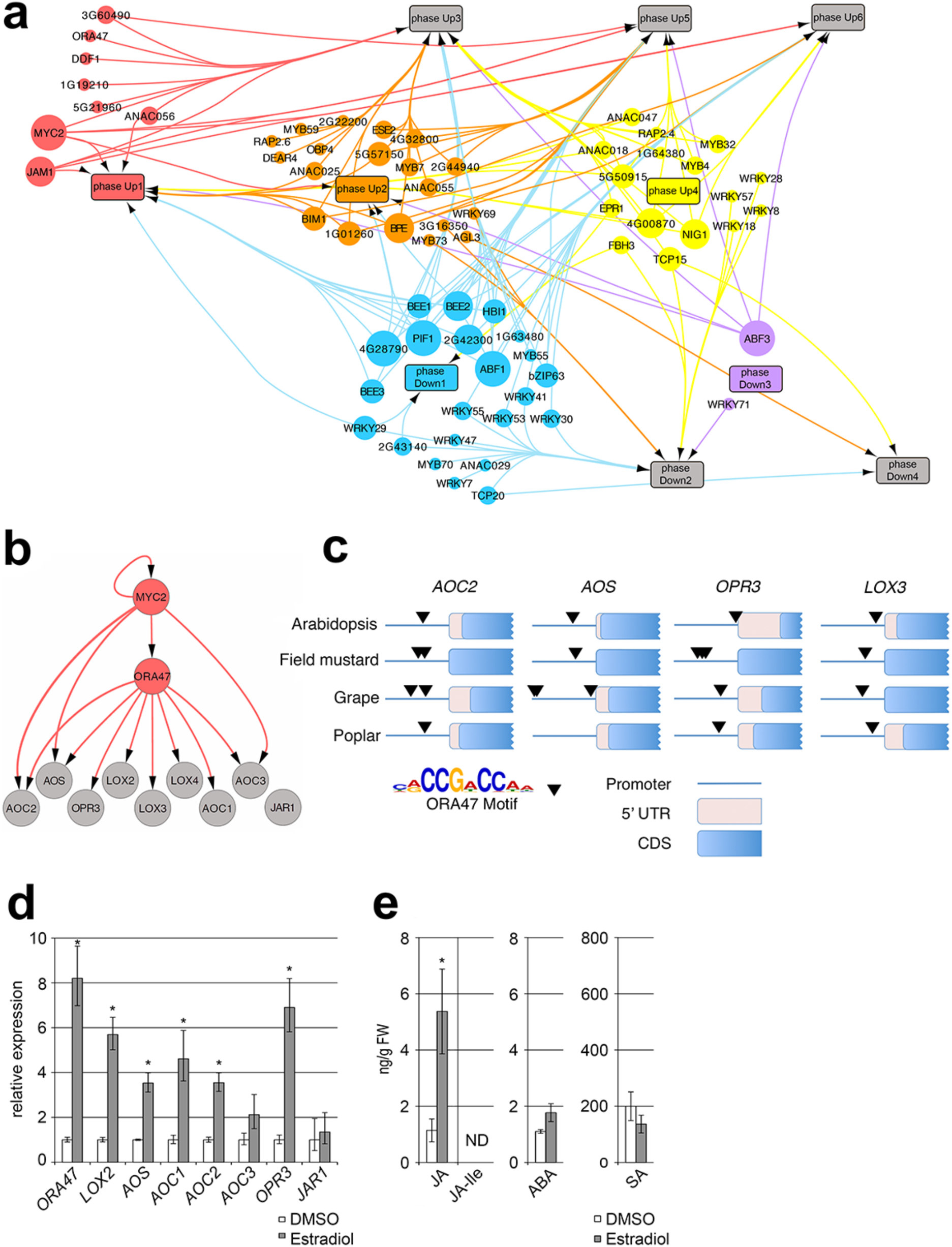
Predicted directional interactions in the JA gene regulatory network and functional analysis of inferred ORA47-controlled subnetwork regulating JA biosynthesis. **(a)** Network plot of inferred connections between MeJA-induced TFs and genes in transcriptional phases. The promoter sequences of genes associated with a transcriptional phase were tested for overrepresentation of DNA motifs shown to be bound to TFs that are differentially transcribed following MeJA treatment. Each TF with a known motif is represented by a colored circle, and is plotted at the time point that its corresponding gene is first differentially expressed. Each transcriptional phase is represented by a rectangle and plotted in time according to its onset. An edge between a TF and a phase indicates significant enrichment of the corresponding binding motif in that phase. The size of each TF node is proportional to the number of phases in which its binding site is overrepresented. To aid interpretation of the network, nodes are grouped and colored according to the transcriptional phase where they first become differentially expressed. (b) Expanded sub-network extracted from the global JA gene regulatory network, indicating regulation of JA biosynthesis genes. Nodes indicating TFs and JA biosynthesis genes are colored grey and orange, respectively. Directed edges (arrows) indicate occurrence of TF-binding sites in the promoter of the target gene. (c) Evolutionary conservation of ORA47 DNA-binding motif. Occurrences of the ORA47 motif (consensus, CCG(A/T)CC) were identified in promoters of an orthologous gene from each of the indicated JA biosynthesis genes (top row). Black arrows indicate a significant match within a gene promoter to the ORA47 motif. 5’UTR, 5-prime untranslated region; CDS, coding sequence. (d) Induction of genes encoding JA biosynthesis enzymes in estradiol-inducible *ORA47* plants. Expression levels of JA biosynthesis genes were measured in leaves 8 h after application of either estradiol or DMSO (mock) using quantitative RT-PCR (qRT-PCR). Shown are the mean expression levels of five biological replicates with mock treatments set at 1. Asterisk indicates significant differences between mock- and estradiol-treated plants (Student’s *t* test; *P* ≤ 0.05; error bars are SE). (e) Production of JA, JA-Ile, ABA, and SA in estradiol-inducible *ORA47* lines. Compound levels were measured from the same leaf tissue harvested for the qRT-PCR analysis described in D. Asterisk indicates significant difference between mock- and estradiol-treated plants (Student’s *t* test; *P* ≤ 0.05; error bars are SE).

Phase Up1 also contains TF genes that are predicted to have a more limited regulatory scope, such as the ERF TF gene *ORA47,* of which the binding motif (consensus, CCG(A/T)CC) is only overrepresented in the promoters of genes assigned to phase Up3. These genes include the JA biosynthesis genes *LOX2, AOS, AOC1,2,3, ACX* and *OPR3,* thus suggesting that this *cis*-element and its cognate TF ORA47 may play a role in regulating JA production, which reflects the positive feedback loop that is known to maintain and boost JA levels upon initiation of the JA response^1^. Focusing on this predicted subnetwork (Figure 5B), we found that *ORA47* and several of the JA biosynthesis genes were predicted to be targets of MYC2, suggesting that MYC2 together with ORA47 regulates JA biosynthesis in *Arabidopsis.* Figure 5C shows that the presence of the ORA47-binding motif is conserved between the promoters of *AOS, AOC2, OPR3* and *LOX3* orthologs of field mustard *(Brassica rapa),* grape *(Vitis vinifera),* and poplar *(Populus trichocarpa*), pointing to a role for ORA47 and its cognate binding element in the regulation of JA biosynthesis genes. Evidence for this is provided by the direct binding of ORA47 to promoter elements of *AOC1, AOC3* and *LOX3,* as demonstrated by yeast one-hybrid experiments (Figure 5-figure supplement 1). Moreover, in stimulated estradiol-inducible *ORA47* plants expression of *LOX2, AOS, AOC1, AOC2* and *OPR3* is increased and accumulation of JA is enhanced (Figure 5D and 5E), which is in line with recent findings^23, 39^. JA-Ile levels did not rise in the *ORA47-* expressing plants, however; this could be explained by our observation that expression of amino acid-conjugating *JAR1*, whose promoter does not contain the ORA47-binding motif, is unchanged. Taken together, these results confirm the predicted role of ORA47 as an important regulator of JA biosynthesis and highlight the potential of combining time series expression data with motif analysis to reveal novel key regulators in gene regulatory networks.

For the vast majority of TFs in our model, it is unclear which specific JA-responsive genes they regulate. To validate and extend our chronological network model further, we made use of transcriptome data sets of two *Arabidopsis* lines that are perturbed in a TF that is predicted by our model to regulate downstream subnetworks. We investigated the effect of the TFs RAP2.6L and ANAC055, which have previously been suggested to regulate JA-responsive genes^40, 41^, by studying their target genes in RAP2.6L-overexpressing and *anac055* mutant *Arabidopsis* lines. Transcriptional profiling of leaves from plants overexpressing *RAP2.6L (RAP2.6L-OX)* under non-stress conditions revealed 93 DEGs (Figure 5-source data 2). Of these, a significant portion of 31 DEGs (P < 3.59e-05; hypergeometric test) was also differentially expressed in the MeJA time series. Projecting the common set of DEGs onto the transcriptional network model revealed that >90% of these genes are present in transcriptional phases that are temporally downstream of the phase containing *RAP2.6L* (phase-Up2, Figure 5-figure supplement 2). Analysis of the overlap between *RAP2.6L*-OX DEGs and the MeJA-induced co-expression clusters from the present study revealed a specific enrichment for RAP2.6L targets in cluster 14, which as described above is itself overrepresented for genes associated with aliphatic glucosinolate production. Interestingly, a recent study showed that RAP2.6L can interact with several aliphatic glucosinolate biosynthetic gene promoters and moreover, that *rap2.6l* mutants were perturbed in glucosinolate production^42^.

Using a similar approach, 56 genes differentially expressed in an *anac055* mutant line compared to wild-type plants (described previously^43^) were overlaid on the JA gene regulatory network. The overlap between MeJA-responsive and ANAC055-regulated genes was statistically significant (24 DEGs, *P* < 4.74e-10; hypergeometric test) and > 85% of these genes became for the first time differentially expressed after *ANAC055* was induced by MeJA (phase-Up2, Figure 5-figure supplement 3). Down-regulated gene co-expression cluster 20 is overrepresented for ANAC055 targets that are enhanced in the *anac055* mutant, and is enriched for GO terms related to SA biosynthesis. Interestingly, ANAC055 has previously been shown to target SA biosynthetic and metabolic genes to negatively regulate SA accumulation following induction by the bacterial toxin coronatine^44^. Collectively, analysis of the transcriptomes of *RAP2.6L-*OX and *anac055* suggests that in the context of the JA gene regulatory network, RAP2.6L and ANAC055 may play a role in the regulation of aliphatic glucosinolate biosynthesis and SA accumulation, respectively. Thus, these two examples demonstrate the value of leveraging TF perturbation data with our information-rich dataset to begin to explore specific transcriptional subnetworks.

### Enhanced induction of dominant-negative splice variant *JAZ10.2* by MeJA

Alternative splicing of mRNA results in the generation of multiple protein isoforms encoded by a single gene. Previously, alternative splicing of *JAZ10* was implicated as a regulatory mechanism of JA-responsive gene expression in *Arabidopsis^9^*. Four distinct splice variants can be produced from the *JAZ10* gene that translate into protein isoforms differing in their sensitivity to COI1-mediated degradation in response to JA accumulation^9^. In particular, the JAZ10.2, 10.3 and 10.4 isoforms lack a functional Jas domain that is important for interaction with COI1, resulting in increased stability and dominant-negative effects on the JA response^9^. Besides JAZ10, JAZ1, 3, 4, 9 and 11 also have annotated mRNA splice variants, but *JAZ3* and *4* are not expressed at a sufficient level in our data set to estimate their relative abundance. While for *JAZ1*, *9* and *11* the relative abundance of the different splice variants did not change over time following application of MeJA (Figure 6-figure supplement 1), the *JAZ10* variants were differentially regulated. Under non-inducing conditions, approximately 60% of *JAZ10* mRNA are of the full-length *JAZ10.1* variant, while the shorter variants *JAZ10.2, 10.3,* and *10.4,* encoding dominant-negative JAZ10 isoforms, each constitute 10-20% of the *JAZ10* mRNA population (Figure 6). Upon exogenous application of MeJA, the fraction of *JAZ10.1* mRNA decreases within 15 min to 13% of the total *JAZ10* mRNA pool. At the same time *JAZ10.2* mRNA (which translates to the same protein sequence as the JAZ10.3 isoform that is proven dominant negative^9^) becomes the most abundant splice variant with 59% of the total *JAZ10* mRNA amount. This change in *JAZ10* splice variant abundance is reverted back to pre-induced ratios within 1.5 h after MeJA treatment. *JAZ10.3* and *JAZ10.4* transcripts remain at a stable level of 10-20% throughout the time course (Figure 6). These results indicate that a temporary shift from predominant *JAZ10* mRNA isoform *JAZ10.1* to *10.2* characterizes the immediate-early response to JA accumulation, which may represent a control mechanism to initially dampen JA-inducible gene expression before JAZ-mediated repression of the JA signaling pathway is released by a relative increase in the JAZ10.1 protein isoform (Figure 6-figure supplement 2).

**Figure 6.**
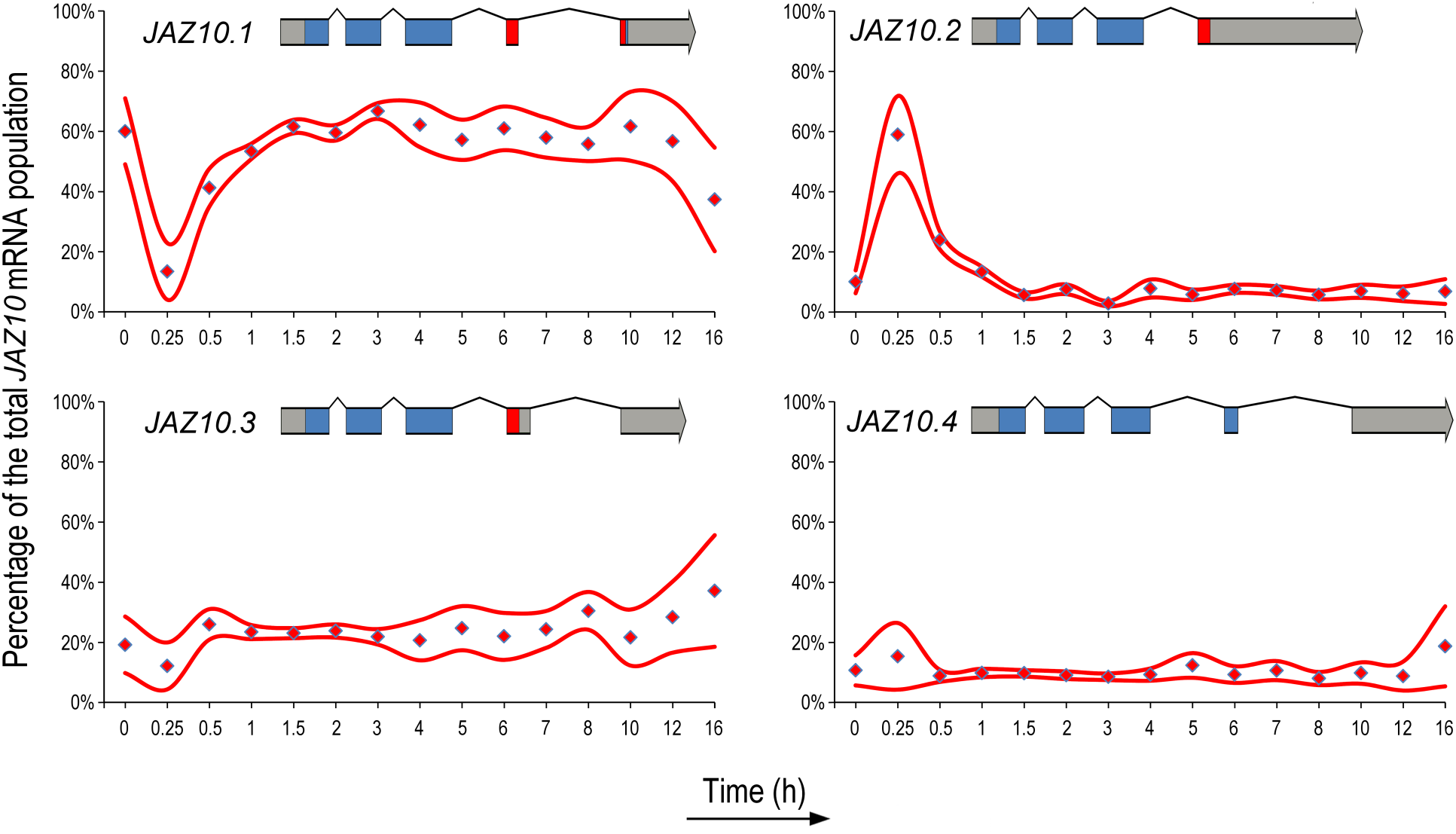
MeJA-induced alternative splicing of *JAZ10* mRNA. The relative abundance of the annotated *JAZ10* transcript splice variants *JAZ10.1, JAZ10.2, JAZ10.3* and *JAZ10.4* after MeJA treatment over time. Each graph indicates the mean percentage (red diamond) of the indicated *JAZ10* splice variant and the 95% confidence interval of four biological replicates (red line) as determined by MISO. In the top of each panel the gene model is depicted; gray (mRNA); blue (CDS) and red (Jas domain).

## DISCUSSION

Computational analyses of high-density time series of RNA-Seq data obtained from Arabidopsis leaves of the same developmental stage (leaf number 6), allowed us to provide an unprecedentedly detailed insight into the architecture and dynamics of the JA gene regulatory network. Previously, studies on phytohormone-induced transcriptional responses have typically included only a limited number of time points or focused on the effect of perturbation of specific regulatory proteins on transcriptional activity in hormone-controlled gene regulatory networks^13, 45^. Our study yielded a considerably more comprehensive MeJA-inducible gene set compared to previous transcriptomic studies (Figure 1). Moreover, it unveiled a variety of detailed temporal expression patterns (Figure 2), including that of transcripts not represented on microarrays and of alternative splice variants (Figure 6), which yielded novel insights into the chronology and regulation of the biologically relevant JA response.

Using a dynamic network approach, we systematically determined how the diverse positive and negative regulatory components in the JA gene regulatory network function over time. MeJA-induced gene activation or repression is shown to be controlled by short transcriptional cascades, yet yielding distinctive transcriptional signatures that correspond to specific sets of genes and biological processes (Figure 2). In general, bHLH TFs are master regulators controlling the majority of the MeJA-inducible genes, while ERF and MYB TFs fine-tune the expression of dedicated sets of target genes (Figure 2, 3 and 4). Besides the known regulators of the JA pathway, several other TFs, whose functions were not previously linked to JA responses, were identified in the network. By using a guilt-by-association approach, twelve early MeJA-induced TFs with unknown roles in the JA response were selected for validation of their biological function in pathogen or insect resistance. Four of these (bHLH27, ERF16, MYB59, and ANAC056) were found to play a role in resistance against the pathogen *B. cinerea* and/or the insect *M. brassicae* (Figure 2), highlighting the high success rate of our approach in the discovery of biological functions of novel genes in the JA network. Numerous other early- and late-expressed TF or enzyme encoding genes still await further exploration for functionality. The wealth of information that is provided by the gene co-expression clusters with their enriched TF-binding motifs and associated biological processes (Figure 3) foretells important regulatory and biological function of individual genes in the JA network.

Our time series data discerned a chronological order of 10 transcriptional phases, showing that the onset of up-regulation preceded that of down-regulation, and that the first phase that was initiated within 15 minutes was represented by transcriptional regulators (Figure 4). JA biosynthesis is shown to be a first target for activation, followed by secondary metabolism. This latter observation correlates with the later activation of many *MYB* TF genes, which are important regulators of secondary metabolism, and the enrichment of MYB DNA-binding motifs in the up-regulated genes in later phases. Down-regulated genes showed enrichment in WRKY TF-binding motifs, which is linked with the suppressed expression of SA-associated defense genes. Integrating TF DNA-binding motif enrichment data with our chronological JA network model predicted putative causal regulations between TFs and downstream JA-regulated subnetworks (Figure 5). Exploring these predictions highlighted a local regulatory module centered around the early JA-responsive ERF TF ORA47. Our *in silico* predictions combined with experimental validation underscore ORA47 as a central regulator of JA biosynthesis, which may form part of a conserved JA amplification loop. For many known and unknown JA-responsive TFs, their exact role in the JA network has remained unclear. Using the established stress-associated TFs RAP2.6L and ANAC055 as examples, the transcriptome data of their respective OX and mutant lines were projected onto our network model. In both cases, this confirmed and extended the predicted regulatory interactions with distinct downstream targets in the JA network model. Specific co-expressed gene clusters in the JA network were shown to be affected in the TF-perturbed lines, highlighting the strength of our clustering analysis for inferring functional regulation mechanisms. A similar transcriptome overlay approach could be used in future studies to further define the roles of other JA-inducible TFs in JA subnetworks.

Finally, we demonstrated an immediate and transient switch of the predominant transcript isoform *JAZ10.1* to *JAZ10.2,* which then becomes the most abundant splice variant of *JAZ10* mRNA and whose corresponding protein product is less prone to degradation via COI1, thus representing a mechanism to attenuate the JA response. A previous study using RT-PCR technology proposed that induction of the dominant negative *JAZ10.3* and *JAZ10.4* splice variants could down tune the JA response^46^. The primers used in that study were unable to distinguish the *JAZ10.2* from the *JAZ10.3* variant and the relative abundance of all *JAZ10* splice variants was not determined over time, so a possible role for enhanced JAZ10.2 induction was overlooked.

In sum, this study provides detailed insight into the dynamics and architecture of the JA gene regulatory network and offers an information-rich data set with a potentially high success rate for the discovery of genes with so-far unknown functions in JA-regulated responses related to plant immunity, growth and development.

## METHODS

### Plant material and growth conditions

Seeds of *Arabidopsis thaliana* Col-0 (wild type), the *myc2,3,4* triple mutant^12^ (At1g32640/At5g46760/At4g17880) and the following T-DNA insertion mutants and transgenic lines (obtained from the Nottingham Arabidopsis Stock Centre): *ofp1* (At5g01840; SALK_111492C), *myb59* (At5g59780; GK-627C09), *anac056* (At3g15510; SALK_137131C), *rap2.6l* (At5g13330; SALK_051006C), *rap2.6* (At1g43160; SAIL_1225G09), *erf16(-1)* (At5g21960; SALK_053563C), *erf16-2* (At5g21960; SALK_096382C), *at1g10586* (At1g10586; SALK_027725C), *bhlh19* (At2g22760; GABI_461E05), *bhlh27(-1)* (At4g29930; SALK_049808C), *bhlh27-2* (At4g29930; SALK_149244C), *bhlh35* (At5g57150; SALK_100300C), *bhlh92* (At5g43650; SALK_033657C), *bhlh113* (At3g19500; GK_892H04), *ora47* (At1g74930; SALK_109440C), *ora59^47^* (At1g06160; GK-061A12.16), and *ORA47* β-estradiol-mducible TRANSPLANTA line^48^ (N2101685) were stratified for 48 h in water at 4°C prior to sowing on river sand. After 2 weeks, the seedlings were transferred to 60-mL pots containing a soil:river sand mixture (12:5) that had been autoclaved twice for 1 h. Plants were cultivated in standardized conditions under a 10-h day (75 μol/m^2^/s^1^) and 14-h night cycle at 21°C and 70% relative humidity. Plants were watered every other day and received half-strength Hoagland nutrient solution containing 10 mM Sequestreen (CIBA-GEIGY GmbH, Frankfurt, Germany) once a week. To minimize within-chamber variation, all the trays, each containing a mixture of plant genotypes or treatments, were randomized throughout the growth chamber once a week. Mutants or treatments were indicated by colored labels of which the code was unknown by the experimenter. T-DNA insertion lines were confirmed homozygous for the T-DNA in the relevant genes with PCR using the gene-specific primers listed in Supplementary file 1E. The *RAP2.6L* overexpressing line *(RAP2.6L-*OX)^40^ and the background accession (WS), were cultivated as described previously^27^.

### MeJA time series setup

Five-week-old Arabidopsis Col-0 plants were treated by dipping the rosette leaves into a mock or MeJA (Duchefa Biochemie BV, Haarlem, The Netherlands) solution. The mock solution contained 0.015% (v/v) Silwet L77 (Van Meeuwen Chemicals BV, Weesp, The Netherlands) and 0.1% ethanol. The MeJA solution contained 0.015% (v/v) Silwet L77 and 0.1 mM MeJA, which was added from a 1,000-fold stock in 96% ethanol. For time series expression analysis, developmental leaf number 6 (counted from oldest to youngest leaf) was harvested from individual plants for each treatment and time point as indicated in Extended Data Table 1. Each individual leaf corresponds to one biological replicate and four biological repeats for all treatments at 15 time points were sequenced (see below). Harvested *Arabidopsis* leaves were snap frozen in liquid nitrogen.

### Induction of the *ORA47* estradiol-inducible line and hormone analysis

Five-week-old *ORA47* inducible overexpression lines were treated by dipping the rosette leaves into a mock or β-estradiol (Sigma-Aldrich, Steinheim, Germany) solution. The mock solution contained 0.015% (v/v) Silwet L77 and 0.1% DMSO. The β-estradiol solution contained 0.015% (v/v) Silwet L77 and 10 μM β-estradiol, which was added from a 1,000-fold stock in DMSO. Hormone analysis was performed as described previously^49^.

### Insect performance and disease bioassays

*Botrytis cinerea* disease resistance was determined essentially as described previously^50^. In brief, *B. cinerea* was grown on half-strength Potato Dextrose Agar (PDA; Difco BD Diagnostics, Franklin Lakes, NJ, USA) plates for 2 weeks at 22°C. Harvested spores were incubated in half-strength Potato Dextrose Broth (PDB; Difco) at a final density of 5 × 10^5^ spores/mL for 2 h prior to inoculation. Five-week-old plants were inoculated by placing a 5-μ droplet of spore suspension onto the leaf surface. Five leaves were inoculated per plant. Plants were maintained under 100% relative humidity with the same temperature and photoperiod conditions. Disease severity was scored 3 days after inoculation in four classes ranging from restricted lesion (<2 mm; class I), non-spreading lesion (2 mm) (class II), spreading lesion (2-4 mm; class III), up to severely spreading lesion (>4 mm; class IV). The distribution of disease categories between genotypes were compared using a Chi-squared test.

*Mamestra brassicae* eggs were obtained from the laboratory of Entomology at Wageningen University where they were reared as described previously^51^. Newly hatched first-instar (L1) larvae were directly placed on *Arabidopsis* leaves using a fine paintbrush. A single L1 *M. brassicae* larva was placed on individual 5-week-old plants (n=30 per mutant line) and its larval fresh weight was determined after 8 days of feeding. To confine the larvae, every plant was placed in a cup that was covered with an insect-proof mesh. Significant differences in larval weight between genotypes were determined using a two-tailed Student’s *t* test.

### High-throughput RNA-sequencing

*Arabidopsis* leaves were homogenized for 2 × 1.5 min using a mixer mill (Retsch, Haan, Germany) set to 30 Hz. Total RNA was extracted using the RNeasy Mini Kit (Qiagen) including a DNaseI treatment step in accordance with manufacturer’s instructions. Quality of RNA was checked by determining the RNA Integrity Number (RIN) using an Agilent 2100 Bioanalyzer and RNA 6000 Nano Chips (Agilent, Santa Clara, United States). For Illumina TruSeq RNA library preparation (see below) only RNA samples with a RIN value of ≥ 9 were used.

RNA-Seq library preparation and sequencing was performed by the UCLA Neuroscience Genomics Core (United States). Sequencing libraries were prepared using the Illumina TruSeq RNA Sample Prep Kit, and sequenced on the Illumina HiSeq2000 platform with read lengths of 50 bases. In total, 12 randomized samples were loaded per lane of a HiSeq2000 V3 flowcell, and each mix of 12 samples was sequenced in 4 different lanes over different flow cells to account for technical variation. A complete scheme of all biological replicates, technical replicates, barcoding used per sample, lane and flow cell usage is provided in Extended Data Table 1. For each of the 15 time points, 4 biological replicates were sequenced in 4 technical replicates, resulting in ∼60 million reads per sample with a read length of 50 bp single end. Complete sequencing setup details can be found in Extended Data Table 1.

Basecalling was performed using the Casava v1.8.2. pipeline with default settings except for the additional argument ‘–use-bases-mask y50,y6n’, to provide an additional Fastq file containing the barcodes for each read in each sample. Sample demultiplexing was performed by uniquely assigning each barcode to sample references, allowing for a maximum of 2 mismatches and only considering barcode nucleotides with a quality score of 28 or greater. The raw RNA-Seq read data are deposited in the Short Read Archive (http://www.ncbi.nlm.nih.gov/sra/) and are accessible through accession number PRJNA224133.

### Processing of RNA-Seq data and analysis of alternative splicing

Read alignment, summarization, normalization and splice variant analysis followed the pipeline described in ref.^28^. Reads were aligned to the *Arabidopsis* genome (TAIR version 10) using TopHat v2.0.4^52^ with the parameter settings: ‘transcriptome-mismatches 3’, ‘N 3’, ‘bowtie1’, ‘no-novel-juncs’, ‘genome-read-mismatches 3’, ‘p 6’, ‘read-mismatches 3’, ‘G’, ‘min-intron-length 40’, ‘max-intron-length 2000’. Aligned reads were summarized over annotated gene models using HTSeq-count v0.5.3p 9^53^ with settings: ‘-stranded no’, ‘-i gene_id’. Sample counts were depth-adjusted using the median-count-ratio method available in the DESeq R package^54^.

Splice variant analysis was performed using MISO^55^; confidence intervals were calculated using Bayesian statistics on all the Monte Carlo simulations of the four biological replicates provided by the MISO^55^ package.

### Differential gene expression analysis and clustering of co-expressed genes

Genes that were significantly differentially expressed after MeJA treatment compared to mock were identified using a generalized linear model (GLM) with a log link function and a negative binomial distribution. Within this model we considered both the time after treatment and the treatment itself as factors. To assess the treatment effect on the total read count for each gene, a saturated model (total counts ∼ treatment + time + treatment:time) was compared to a reduced model considering time alone (total counts ∼ time) using ANOVA with a Chi-squared test. The obtained *P* values for all genes were corrected for multiple testing using a Bonferroni correction. All genes that did not meet the following requirement were omitted from further analysis: a minimum 2-fold difference in expression on at least one of the 15 time points, supported by a minimum of 10 counts in the lowest expressed sample, and a *P* value ≤ 0.01 for that time point. Remaining genes with Bonferroni-corrected *P* value ≤ 0.05 were called as differentially expressed genes (DEGs). All statistics associated with testing for differential gene expression were performed with R (http://www.rproject.org).

Of all the DEGs, the time point of first differential expression was predicted. To this end the significance of the treatment effect at each time point was obtained from the GLM, represented by its *z* score. These values were used as a basis to interpolate the significance of the treatment effect in between the sampled time points. This was done using the interpSpline function in R using 249 segments. The first time point of differential expression was set where the z score was higher than 2.576 (equivalent of *P* value 0.01) for up-regulation or lower than −2.576 for down-regulation.

Temporal expression profiles of the DEGs were clustered using SplineCluster^56^. Clustering was performed using the profiles of log_2_-fold changes at each time point (MeJA-treated versus mock), with a prior precision value of 10^−4^, the default normalization procedure and cluster reallocation step^57^. All other optional parameters remained as default.

### TF family and promoter motif analysis

To determine which TF families are enriched among the genes differentially expressed in response to application of MeJA, we tested for overrepresentation of 58 TF families described in the TF database PlantTFDB version 3.0^58^. Overrepresented TF families within a set of genes were analyzed using the cumulative hypergeometric distribution, with the total number of protein coding genes (TAIR version 10) as the background. *P* values were corrected for multiple testing with the Bonferroni method.

For promoter motif analysis, the promoter sequences defined as the 500 bp upstream of the predicted transcription start site (TSS) were retrieved from TAIR (version 10). *De novo* promoter motifs were identified by applying the motif-finding programs MEME^59^ and XXmotif^60^ to the promoters of all genes present in a given coexpression cluster. This approach exploits the strengths of different motif-finding strategies, which has been demonstrated to improve the quality of motif detection^61^. Both algorithms searched for motifs on the forward and reverse strands and used the zero-or-one occurrences per sequence (ZOOPS) motif distribution model. MEME was run using a 3rd-order Markov model learned from the promoter sequences of all genes in the *Arabidopsis* genome, using parameter settings: ‘-minw 8 -maxw 12 -nmotifs 10’. XXmotif was run using a 3rd-order Markov model and the medium similarity threshold for merging motifs, with all other parameters kept as default. This analysis yielded a large number of motifs, many of which were highly similar. To reduce redundancy amongst motifs, a postprocessing step was performed using the TAMO software package^62^. Motifs were converted to TAMO format, clustered using the UPGMA algorithm, and merged to produce consensus motifs. The set of processed motifs were converted to MEME format for all subsequent analyses using the tamo2meme function available in the MEME Suite^63^. For the analysis of known motifs originating from protein-binding microarray (PBM) studies^31, 32^, the published weight matrices were converted into MEME format.

The presence or absence of a given motif within a promoter was determined using FIMO^64^. A promoter was considered to contain a motif if it had at least one match with a *P* value ≤ 10^−4^. For each *de novo-* and PBM-derived motif, the statistical enrichment of each motif within the promoters of co-expression gene clusters or transcriptional phases was tested using the cumulative hypergeometric distribution. This test computes the probability that a motif is present within a set of promoter sequences at a frequency greater than would be expected if the promoters were selected at random from the *Arabidopsis* genome.

Analysis of the *ORA47* DNA-binding motif conservation across different plant species was performed using the promoters of genes orthologous to *Arabidopsis AOC2, AOS, OPR3* and *LOX3.* Orthologs were identified in *Vitis vinifera, Populus trichocarpa* and *Brassica rapa* genomes (Ensembl database release 25) using the reciprocal best BLAST hit method^65^. Presence or absence of the *ORA47* motif in the promoters (500 bp upstream of predicted TSS) of these orthologous genes was determined using FIMO as described above.

### Gene Ontology analysis

Gene ontology (GO) enrichment analysis on gene clusters was performed using GO term finder^66^ and an *Arabidopsis* gene association file downloaded from ftp.geneontology.org on 2nd May 2013. Overrepresentation for the GO categories ‘Biological Process’ and ‘Molecular Function’ were identified by computing a *P* value using the hypergeometric distribution and false discovery rate for multiple testing (*P* ≤ 0.05).

### Identification of chronological phases in MeJA-induced gene expression

To identify phases of MeJA-induced changes in transcription we first divided all DEGs depending on whether they were either up- or down-regulated in response to MeJA and then further according to their function as either a transcriptional regulator (termed regulator genes) or having a different function (termed regulated genes). To identify DEGs that encode transcriptional regulators we used the comprehensive list of Arabidopsis TFs and transcriptional regulators described in ref.^67^ and subjected it to minor additional manual literature curation. This filtering yielded four mutually exclusive sets of MeJA-responsive genes (i.e. regulator genes up and down, regulated genes up and down). For each of the four gene sets, the depth-normalized expression values (see above) for all pairs of time points were compared pairwise using the Pearson correlation measure. Each resulting correlation matrix was then clustered using the Euclidean distance measure with average linkage. The resulting dendrograms were used to infer distinct phases of MeJA-induced transcription, where each phase has a start and end time. Each gene present in one of the four final gene sets was assigned to a transcriptional phase based on its time point of first differential expression (Figure 4-figure supplement 1). All genes that were for the first time differentially expressed before, or equal to, the final time point in a given phase (clustered group of time points), and after the final time point of a preceding phase, were assigned to that transcriptional phase (see Figure 4-figure supplement 2 for overview of the method).

### Network construction

The identification of potential regulatory network connections between TFs and transcriptional phases was performed with a set of TFs that met two criteria: (1) They were differentially expressed in response to application of MeJA (and thus belong to a phase). (2) They have an annotated DNA-binding motif (as described in “Promoter motif analysis”). Each set of genes that constitute a transcriptional phase (10 phases in total) was tested for overrepresentation of each motif using the hypergeometric distribution as described above. A directional edge was drawn from a TF to a phase when its cognate binding motif was overrepresented in the promoters of genes belonging to that phase (hypergeometric distribution; *P* ≤ 0.005). The resulting network was visualized using Cytoscape^68^.

### Quantitative RT-PCR analysis

For quantitative RT-PCR (qRT-PCR), RNA was extracted as described in ref.^69^ and subsequently treated with DNaseI (Fermentas, St. Leon-Rot, Germany) to remove genomic DNA. Genomic DNA-free total RNA was reverse transcribed by using RevertAid H minus Reverse Transcriptase (Fermentas, St. Leon-Rot, Germany). PCR reactions were performed in optical 384-well plates with a ViiA 7 realtime PCR system (Applied Biosystems, Carlsbad, CA, USA), with SYBR^®^ Green (Applied Biosystems, Carlsbad, CA, USA). A standard thermal profile was used: 50°C for 2 min, 95°C for 10 min, followed by 40 cycles of 95°C for 15 s and 60°C for 1 min. Amplicon dissociation curves were recorded after cycle 40 by heating from 60 to 95°C with a ramp speed of 0.05°C/sec. All primers used for qRT-PCR are listed in Supplementary file 1E. The gene *At1g13320* was used as reference for normalization of expression^70^.

### Microarray analysis of *RAP2.6L* transgenic plants

Total RNA was extracted from three leaves per plant (28-days-old), labeled and hybridized to CATMA v4 arrays^71^ as described previously^25^. Three biological replicates of WS and *RAP2.6L-*OX samples were pooled separately and labeled three times with each dye to give six technical replicates. Analysis of expression differences between WS and *RAP2.6L-*OX was performed with the R Bioconductor package limmaGUI^72^ using Print-Tip lowess transformation and quantile-normalization.

### Yeast-1-Hybrid (Y1H) protein-DNA interaction assays

Cloning of bait promoter DNA and yeast transformation was performed as described in ref.^43^. All primers that were used to clone promoter fragments are listed in Extended Data Table 13. *ORA47* coding sequence was isolated from the transcription factor library described in ref.^43^ and the correct sequence confirmed by sequencing (Supplementary file 1E). Prey strains were constructed by cloning the *ORA47* coding sequence into pDEST22 (Invitrogen) and transforming AH109 yeast (Clontech), while empty pDEST22 was used to transform AH109 as a negative control. Three μL of bait strain cultures were spotted onto YPDA (yeast, peptone, dextrose, adenine) plates and dried before being overlaid with 3 μL of prey strain culture and left to grow overnight at 30^o^C. Colonies were subcultured in 1 mL mating selective media (SD-Leu-Trp, Clontech) and grown for two nights at 30^o^C with shaking. Cultures were diluted to 10^8^ cells/mL in SD-Leu-Trp liquid media before four 10-fold serial dilutions were made. Three μL of each diploid strain was plated to mating selective (SD-Leu-Trp, Clontech) and interaction selective (SD-Leu-Trp-His, Clontech) media and incubated at 30^o^C for 72 h before being photographed using a G:Box EF2 (Syngene). For promoter D, 5 mM 3-Aminotriazole (Sigma-Aldrich) was required to suppress autoactivation of *HIS3* expression by this promoter region. For promoters A, B and D experiments were performed using two independent promoter transformants and four transcription factor transformants, for a total of eight replicates. For promoter C, there were three replicates across two independent promoter transformants and two transcription factor transformants.

## Acknowledgments

We would like to thank Leon Westerd from the laboratory of Entomology at Wageningen University for providing *Mamestra brassicae* eggs. The *RAP2.6L-*OX line and background accession (WS) were obtained from Nataraj Kav (University of Alberta, Edmonton, Canada). This work was supported by the Netherlands Organization for Scientific Research (NWO) through the Dutch Technology Foundation (STW) VIDI Grant no. 11281 (to SCMvW), STW VENI Grant no. 13682 (to RH), Biotechnology and Biological Sciences Research Council (BBSRC) Grants BB/F005806/1 (to KD) and BB/M017877/1 (to KD and AT), and ERC Advanced Grant no. 269072 (to CMJP) of the European Research Council. Engineering and Physical Sciences Research Council/BBSRC-funded Warwick Systems Biology Doctoral Training Centre (AJ), and BBSRC Systems Approaches to Biological Research studentship (JR).

## Author contributions

SCMvW and CMJP conceived the approach and together with MCvV and RJH designed the study. MCvV and RJH designed all bioinformatics approaches. KD provided analytical and intellectual contributions. MCvV and RJH performed data analysis. MCvV, RJH, AJHvD, MPM, IAV, LC, MS, GJW and IvdN performed mutant genotyping and validation experiments. MdV and RCS performed hormone measurements. AJ and AT performed Y1H assays. JR performed microarray experiments. MCvV, RJH, CMJP and SCMvW wrote the manuscript. All authors discussed the results and commented on the manuscript.

---

**Figure 2-figure supplement 1.**
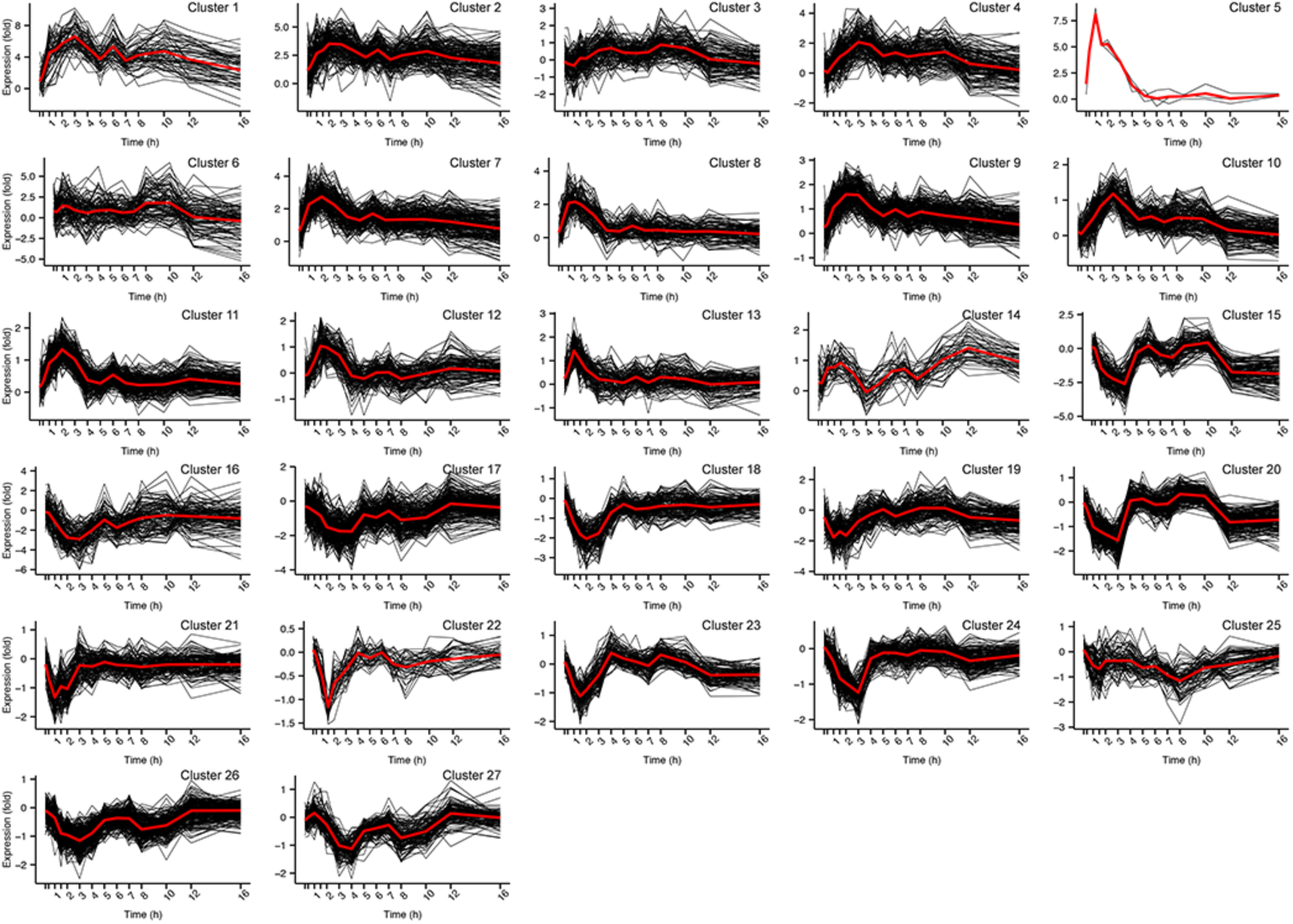
SplineCluster analysis of MeJA-responsive gene expression profiles. Shown are the 27 clusters identified by SplineCluster. Individual genes are plotted in black. Red lines indicate the mean expression level (log2-fold change (MeJA/mock)) for all genes in a cluster.

**Figure 2-figure supplement 2.**
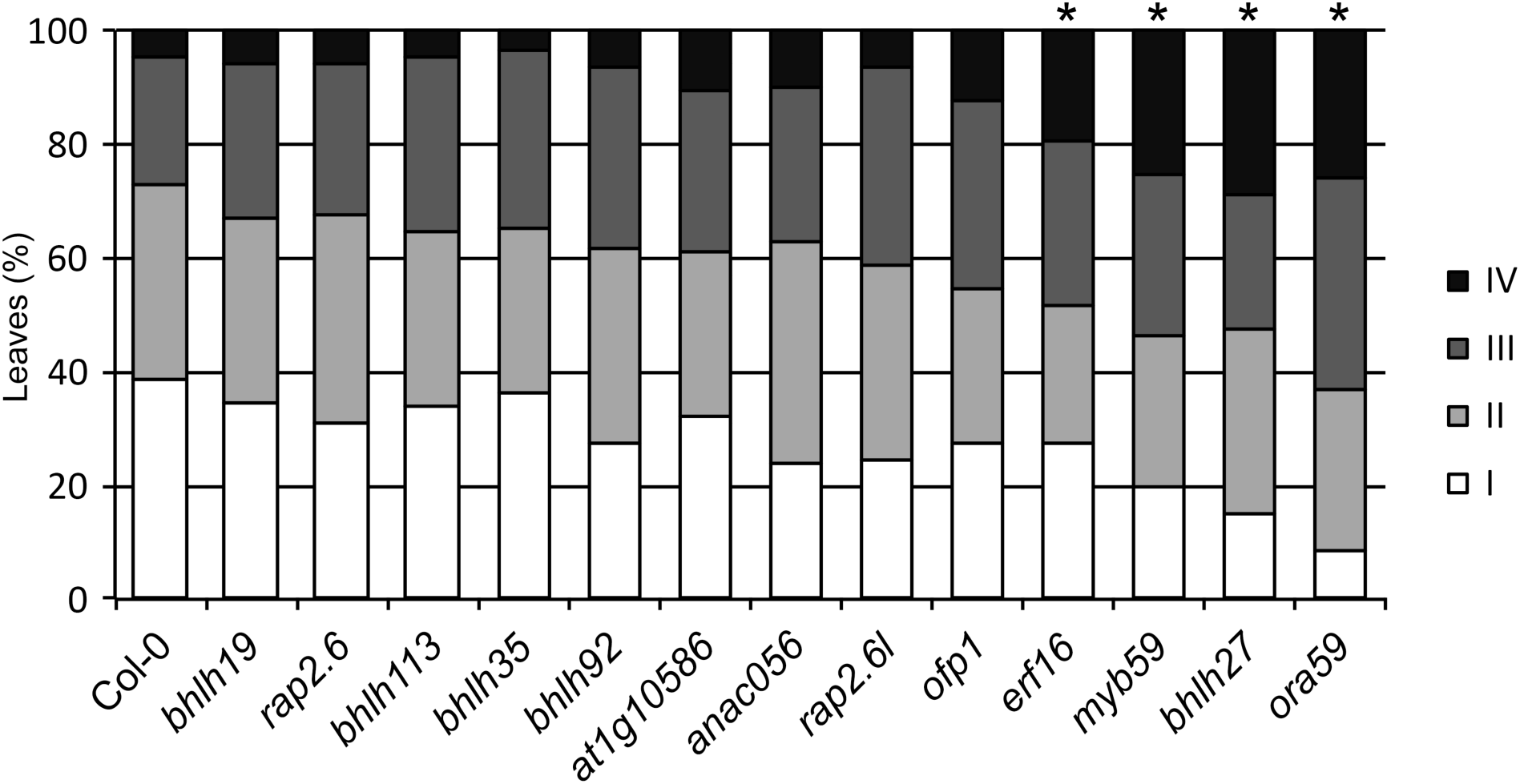
*B. cinerea* disease severity assay with selected mutant lines. Quantification of *B. cinerea* disease severity at 3 days after inoculation of T-DNA insertion lines for selected genes encoding predicted regulators of the JA pathway. Disease severity of inoculated leaves was scored in four classes ranging from restricted lesion (class I), nonspreading lesion (class II), spreading lesion (class III), up to severely spreading lesion (class IV). The percentage of leaves in each class was calculated per plant (n=20). Asterisk indicates statistically significant difference from Col-0 (Chi-squared test; *P* ≤ 0.05). Most genotypes were tested multiple times.

**Figure 2-figure supplement 3.**
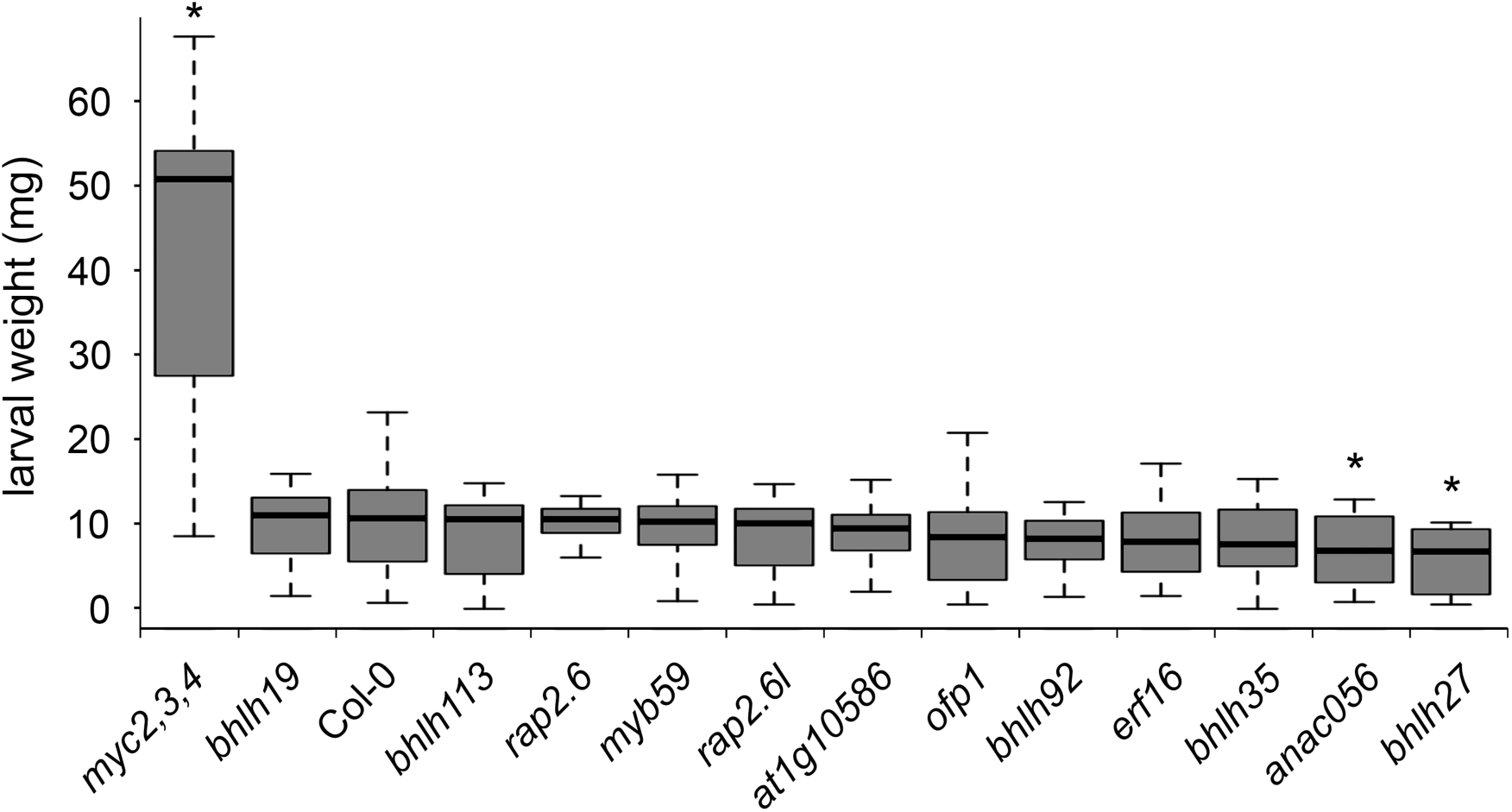
Growth of *M. brassicae* larvae on selected mutant lines. Larval fresh weight was determined after 8 days of feeding on T-DNA insertion lines for selected genes encoding predicted regulators of the JA pathway. Values represent mean weight (±SE) of the larvae. Asterisk indicates statistically significant difference from Col-0 (two-tailed Student’s *t* test for pairwise comparisons; *P* ≤ 0.05; n=30; error bars are SE). Most genotypes were tested multiple times.

**Figure 2-figure supplement 4.**
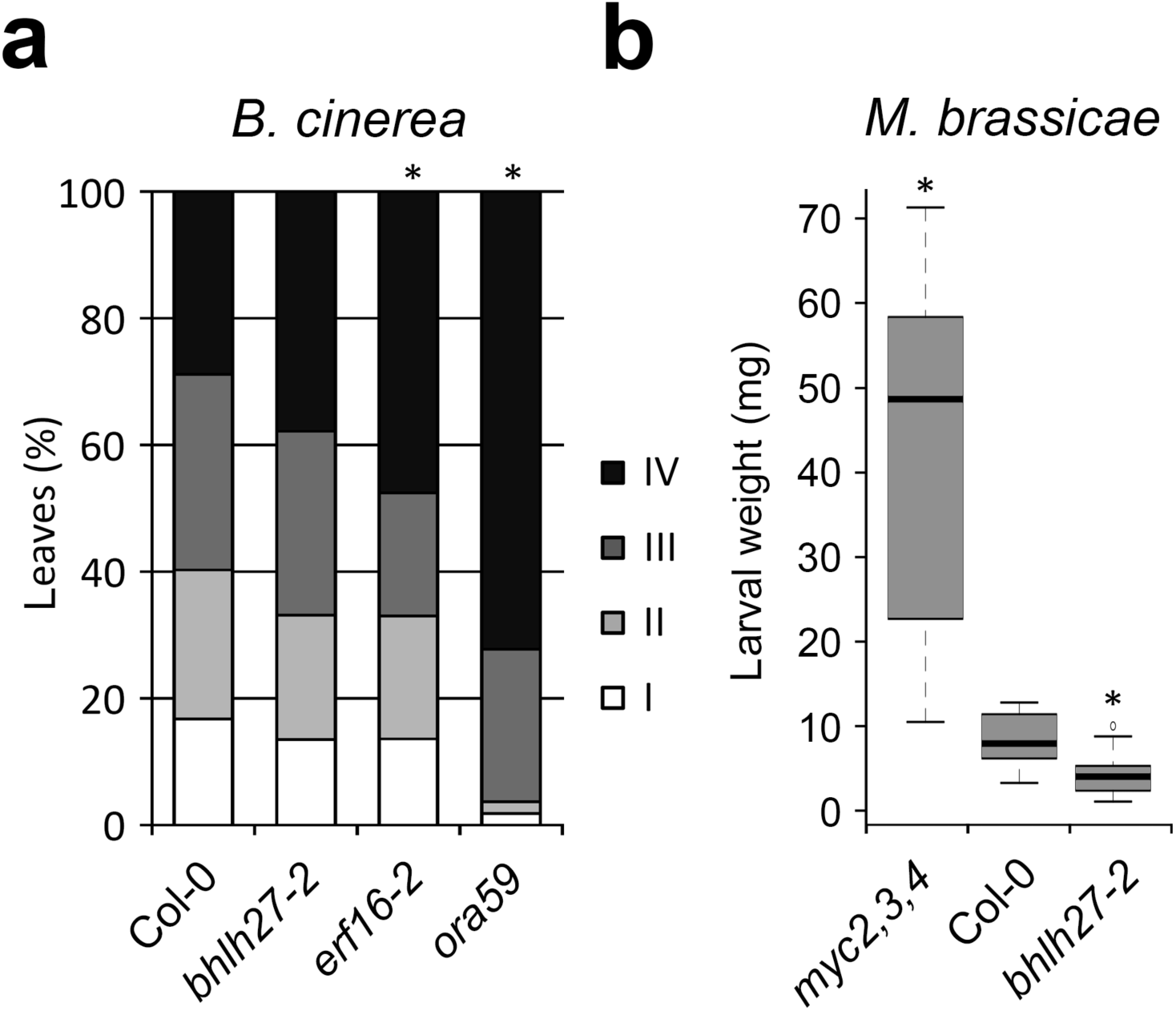
*B. cinerea* disease severity and growth of *M. brassicae* larvae on additional mutant alleles. **(a)** Quantification of *B*. *cinerea* disease severity at 3 days after inoculation of T-DNA insertion lines harboring an *bHLH27* or *ERF16* mutation *(bhlh27-2, erf16-2)* different from that in the mutant lines tested in main Figure 2 (and supplements 2 and 3). Disease severity of inoculated leaves was scored in four classes ranging from restricted lesion (class I), nonspreading lesion (class II), spreading lesion (class III), up to severely spreading lesion (class IV). The percentage of leaves in each class was calculated per plant (n=20). Asterisk indicates statistically significant difference from Col-0 (Chi-squared test; *P* ≤ 0.05). **(b)** Larval fresh weight of *M. brassicae* was determined after 8 days of feeding on *bhlh27-2*. Asterisk indicates statistically significant difference from Col-0 (two-tailed Student’s *t* test for pairwise comparisons; *P* ≤ 0.05; n=>10; error bars are SE).

**Figure 4-figure supplement 1.**
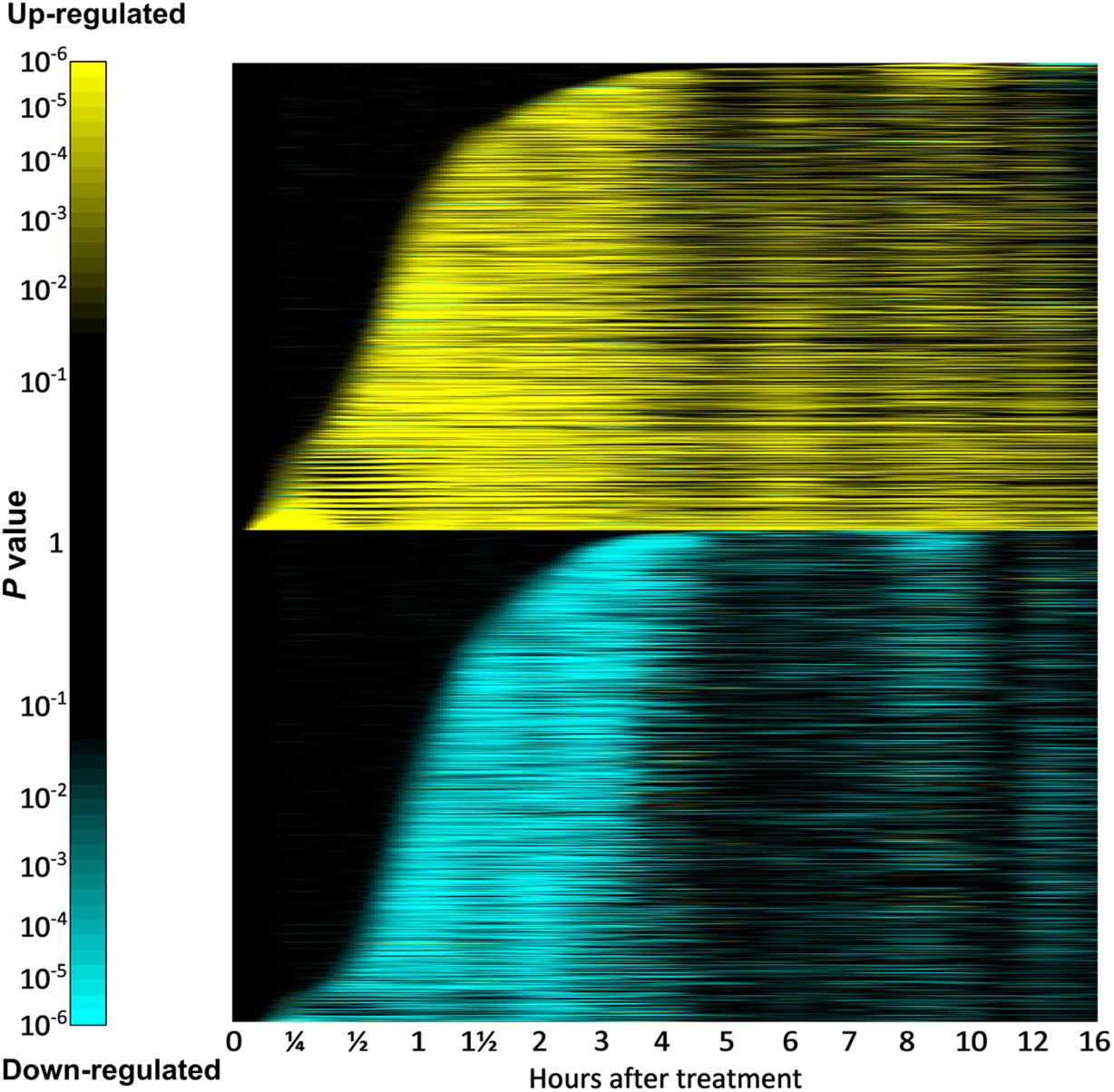
Timing of differential expression for all differentially expressed genes. All differentially expressed genes were divided in two groups, dependent on whether they were up- or down-regulated over time by MeJA treatment. Estimated *z* scores reflect significance of differential expression over time and were used to order genes according to their time of first differential expression. Shown are z scores converted to *P* values.

**Figure 4-figure supplement 2.**
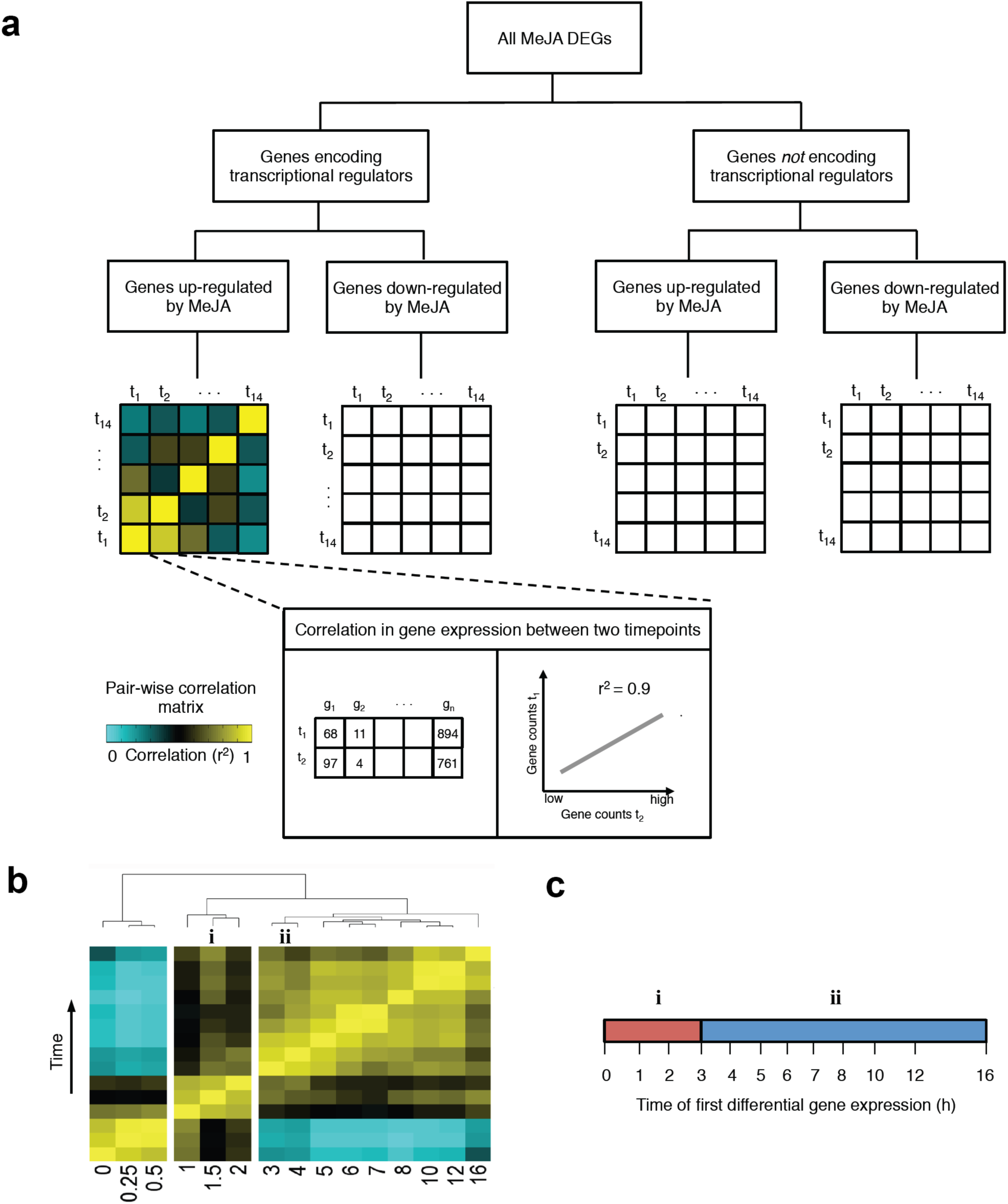
Identification of transcriptional phases induced in response to MeJA treatment. **(a)** Generating correlation matrices for different sets of MeJA-induced genes. The complete list of MeJA-responsive DEGs is progressively filtered into four sets of genes according to two binary criteria: 1) Gene encodes a transcriptional regulator (true or false); 2) Direction of change in expression (up-regulated or down-regulated). For each of the four mutually exclusive gene sets, a correlation matrix of gene transcription counts between all pairs of time points is computed. Inset, an example illustrating how the squared Pearson correlation coefficient (r^2^) is computed between one pair of time points. **(b)** The dendrogram obtained by hierarchical clustering of the transcriptome correlation matrix of each of the four gene sets reveals that there are groups of time points showing highly correlated levels of gene expression within that group, but displaying reduced correlation levels with the time points belonging to other groups. A relatively weak correlation between a pair of adjacent time points signifies a coordinated switch in transcriptional activity of a fraction of the genes. In this example, two phases are identified; the first begins at 1 h (i) and the second at 3 h (ii). **(c)** Each gene present in one of the four final gene sets is assigned to a transcriptional phase based on its time point of first differential expression (Figure 4-figure supplement 2). All genes that are for the first time differentially expressed before, or equal to, the final time point in a given phase (clustered group of time points), and after the final time point of a preceding phase, are assigned to that transcriptional phase. Moreover, if a group of clustered time points includes the 0 time point, then all genes assigned to this group are combined with the genes of the adjacent group of time points, which is numbered as the first transcriptional phase. Red and blue bars indicate time intervals used for assignment of genes to phases i and ii, respectively.

**Figure 5-figure supplement 1.**
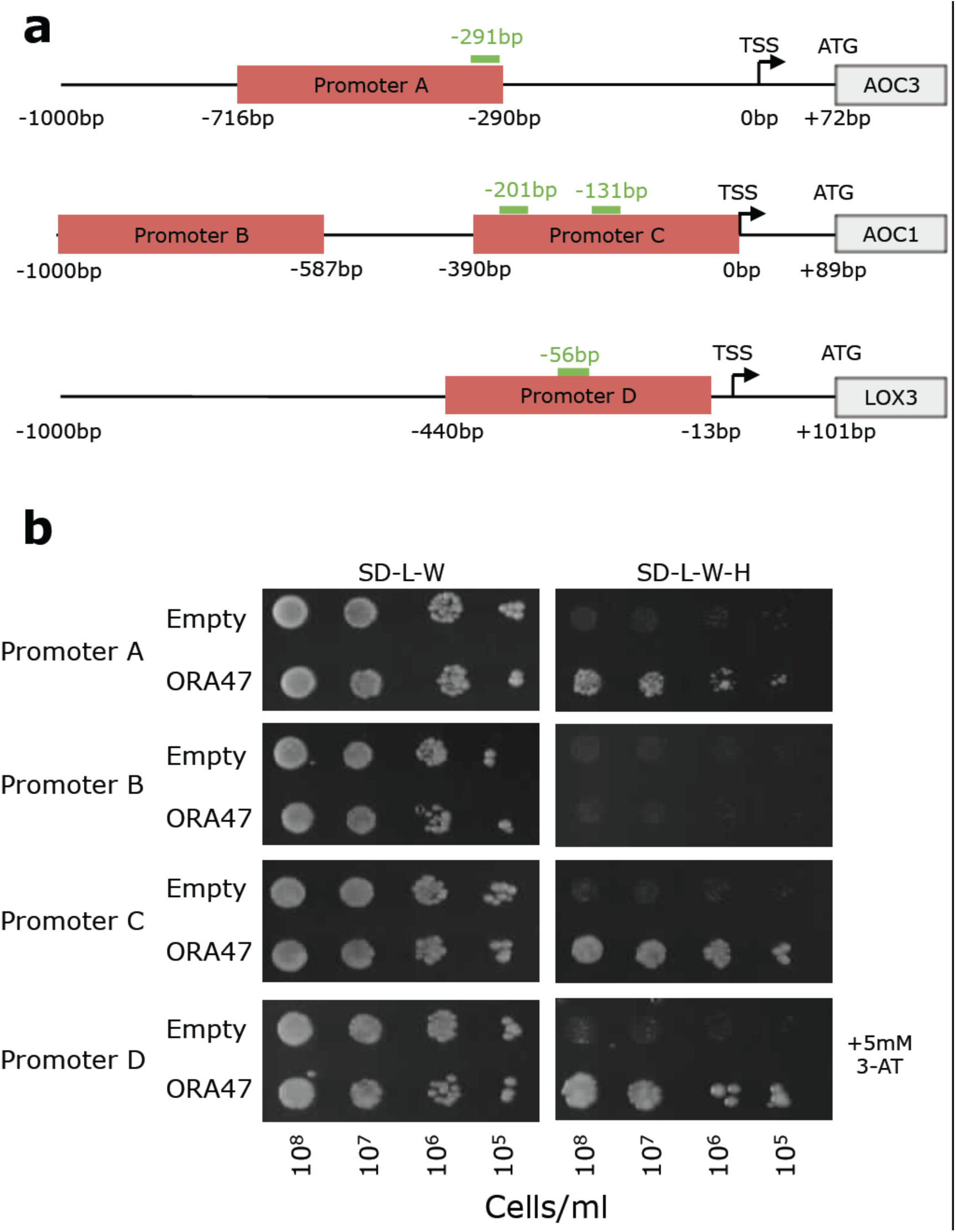
ORA47 can bind to the promoters of multiple *Arabidopsis* genes encoding JA biosynthesis enzymes in yeast. **(a)** Schematic of promoters for JA biosynthesis genes *AOC1, AOC3* and *LOX3.* The locations of fragments used for Y1H assays are shown in red. Occurrences of ORA47 DNA-binding motif are shown in green. Numbers are relative to transcription start site (TSS) **(b)** Interaction between ORA47 and promoter fragments A, C and D was confirmed by growth on SD-Leu-Trp-His (SD-L-W-H) media. Yeast transformed with plasmids containing promoter construct D were grown on SD-L-W-H supplemented with 5 mM 3-AT. Yeast transformed with promoter fragment B (which does not contain a significant match for the ORA47 motif) did not grow on selective media, indicating no interaction between ORA47 and this DNA sequence.

**Figure 5-figure supplement 2.**
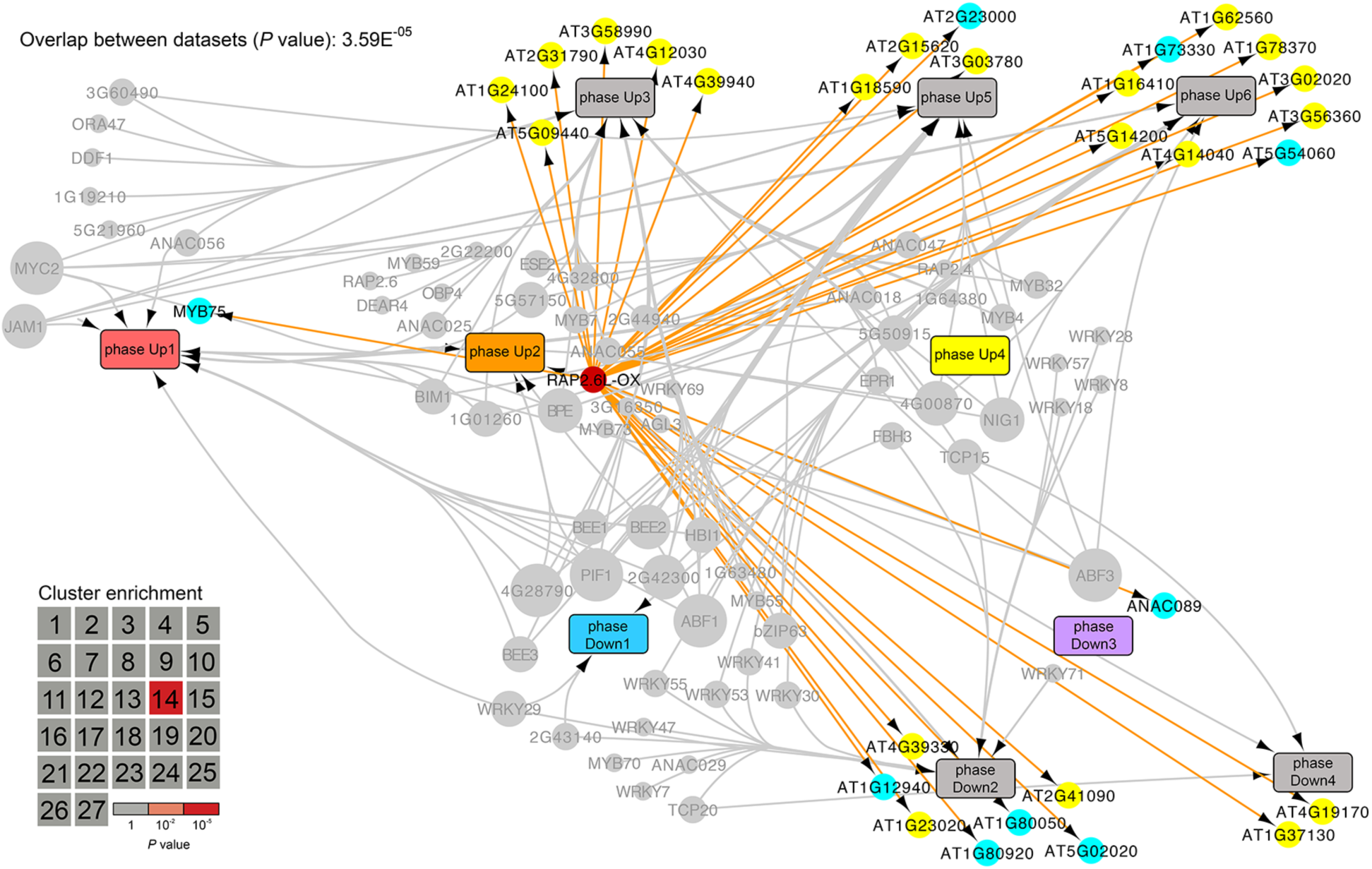
Projection of RAP2.6L target genes on the JA network model. Genes that are differentially expressed in the *RAP2.6L*-OX line were overlaid on the network described in Figure 5. DEGs are indicated by nodes and positioned according to phase membership. Direction of misregulation in *RAP2.6L*-OX is indicated by color; yellow, up-regulated; cyan, down-regulated. The gene encoding RAP2.6L is shown as a red-colored node. Inset: heatmap indicating hypergeometric enrichment *P* value of RAP2.6L target genes in each co-expression cluster.

**Figure 5-figure supplement 3.**
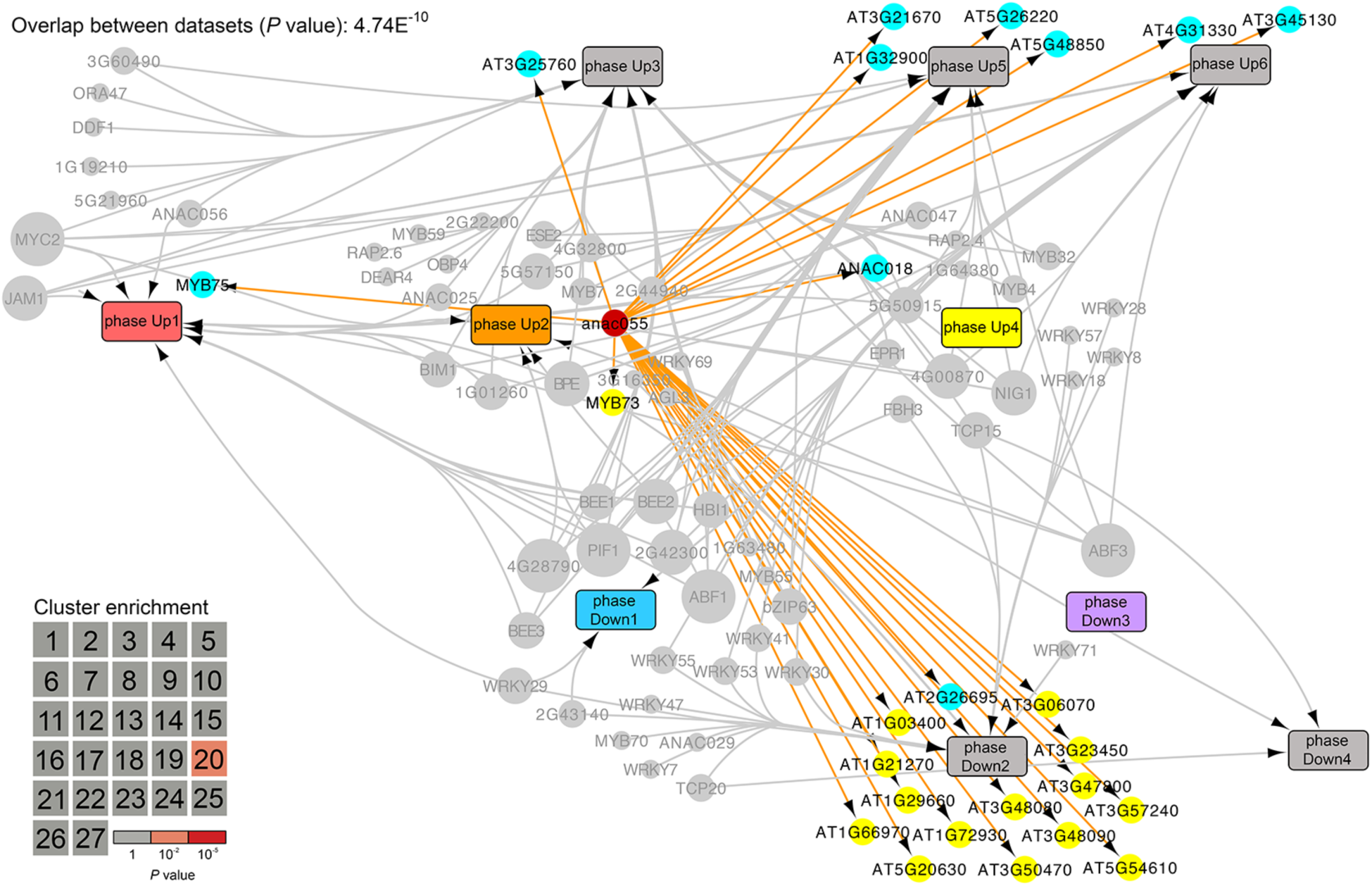
Projection of ANAC055 target genes on the JA network model. Genes that are differentially expressed in the *anac055* mutant were overlaid on the network described in Figure 5. DEGs are indicated by nodes and positioned according to phase membership. Direction of misregulation in the *anac055* mutant is indicated by color; yellow, up-regulated; cyan, down-regulated. The gene encoding ANAC055 is shown as a red-colored node. Inset: heatmap indicating hypergeometric enrichment *P* value of ANAC055 target genes in each co-expression cluster.

**Figure 6-figure supplement 1.**
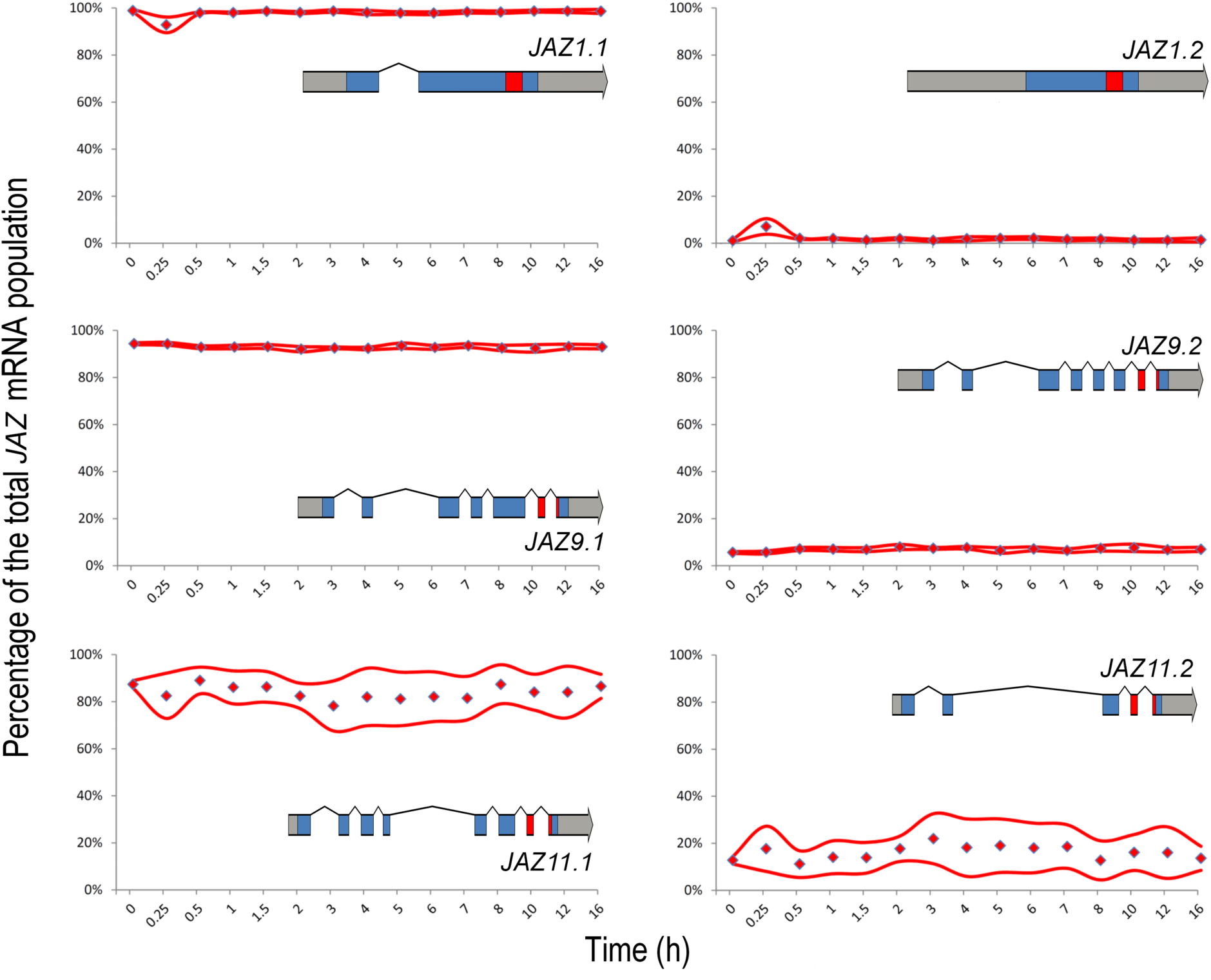
Alternative splicing of *JAZ* transcripts upon MeJA treatment. The relative abundance of each isoform for *JAZ1, JAZ9* and *JAZ11* at each time point was determined using MISO. Each panel depicts the percentage of the total mRNA population that is represented by the indicated isoform for each time point. The red line represents the 95% confidence interval of four biological replicates, derived from MISO. In each panel the representative gene model is depicted; grey (mRNA), blue (CDS) and red (Jas domain).

**Figure 6-figure supplement 2.**
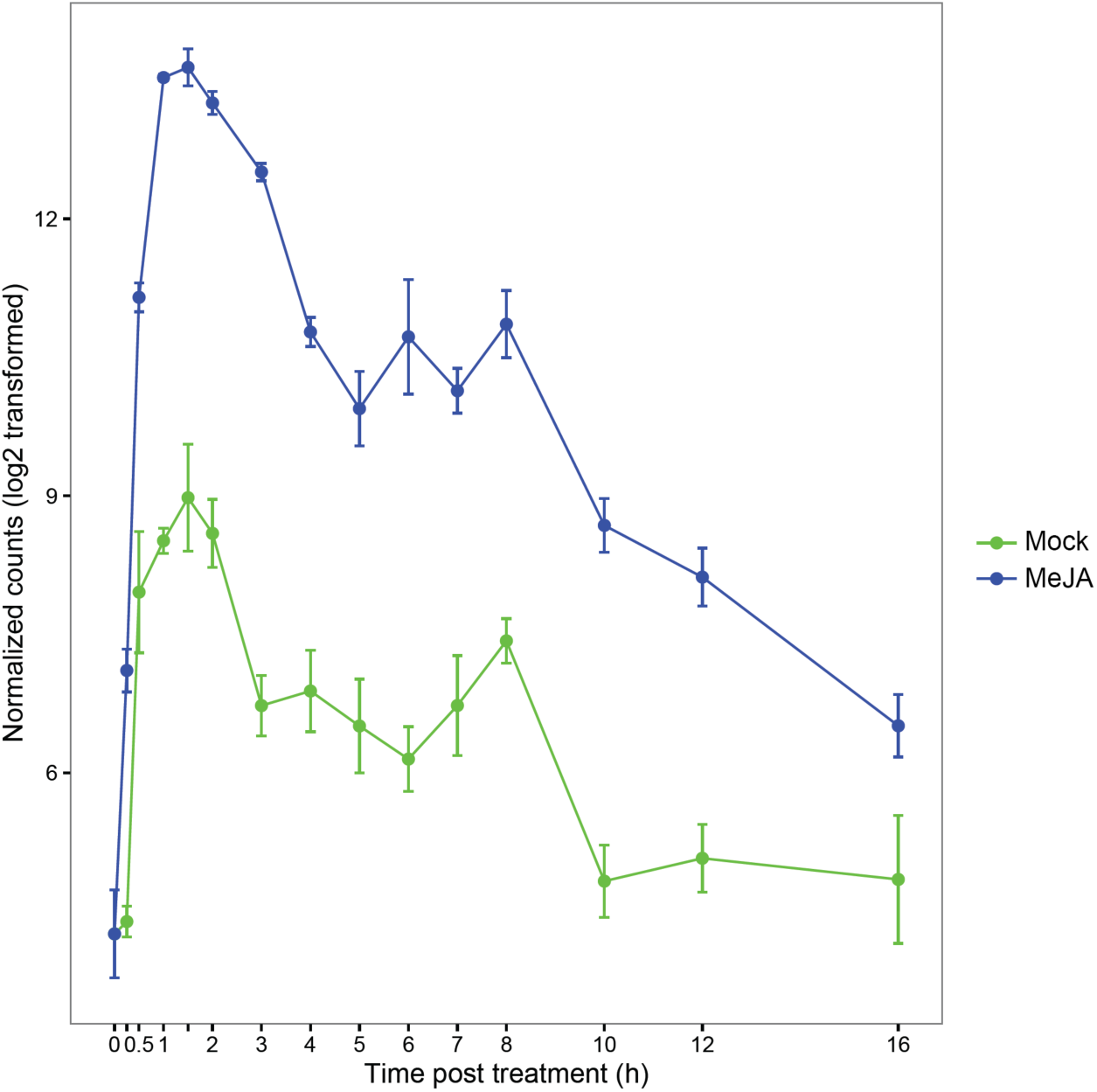
Relative expression of *JAZ10* transcripts upon MeJA treatment. The relative expression of *JAZ10* at each time point is shown after Mock (green) and MeJA treatment (blue). The error bars represent the SE of four biological replicates.

**Figure 2-source data 1** *Arabidopsis* Gene Identifier (AGI) codes for members of each of the 27 gene co-expression clusters identified by SplineCluster.

**Figure 2-source data 2** GO-terms overrepresented in each of the 27 co-expression gene clusters.

**Figure 3-source data 1** Enrichment of known TF DNA-binding motifs in each of the 27 co-expression gene clusters.

**Figure 3-source data 2** *De novo*-derived motif enrichment in each of the 27 gene co-expression clusters.

**Figure 3-source data 3** *De novo*-derived sequence motifs in Weblogo and Position Weight matrix (MEME) format.

**Figure 4-source data 1** *Arabidopsis* Gene Identifier (AGI) codes for members of each of the 10 transcriptional phases that are initiated after MeJA treatment.

**Figure 4-source data 2** GO-terms overrepresented in each of the 10 transcriptional phases that are initiated after MeJA treatment.

**Figure 4-source data 3** Known TF DNA-binding motif enrichment in each of the 10 transcriptional phases that are initiated after MeJA treatment.

**Figure 4-source data 4** *De novo*-derived motif enrichment in each of the 10 transcriptional phases that are initiated after MeJA treatment.

**Figure 5-source data 1** List of differentially expressed TF genes and enrichment of their corresponding TF DNA-binding motif in the promoters of genes within a transcriptional phase.

**Figure 5-source data 2** List of differentially expressed genes obtained from microarray analysis of *RAP2.6L-*OX.

**Supplementary file 1A** Time series experimental set-up and mRNA sequencing details.

**Supplementary file 1B** Median-count ratio normalized expression values of all genes and biological replicates at the 15 time points after MeJA and mock treatments.

**Supplementary file 1C** Mean expression values for all genes across the time series following MeJA treatment. Expression values are represented as fold changes (MeJA/mock) at each time point.

**Supplementary file 1D Significance of differential expression over time for each of the 27 clusters of co-expressed genes in response to MeJA treatment.** For each SplineCluster of co-expressed genes, the estimated z scores were computed per gene using a generalized linear model as described in the materials and methods. The obtained values reflect the significance of differential expression per time point for a given gene. Scores for all genes belonging to a cluster were assembled into a matrix and were hierarchically clustered using the Euclidean distance measure with average linkage.

**Supplementary file 1E.**
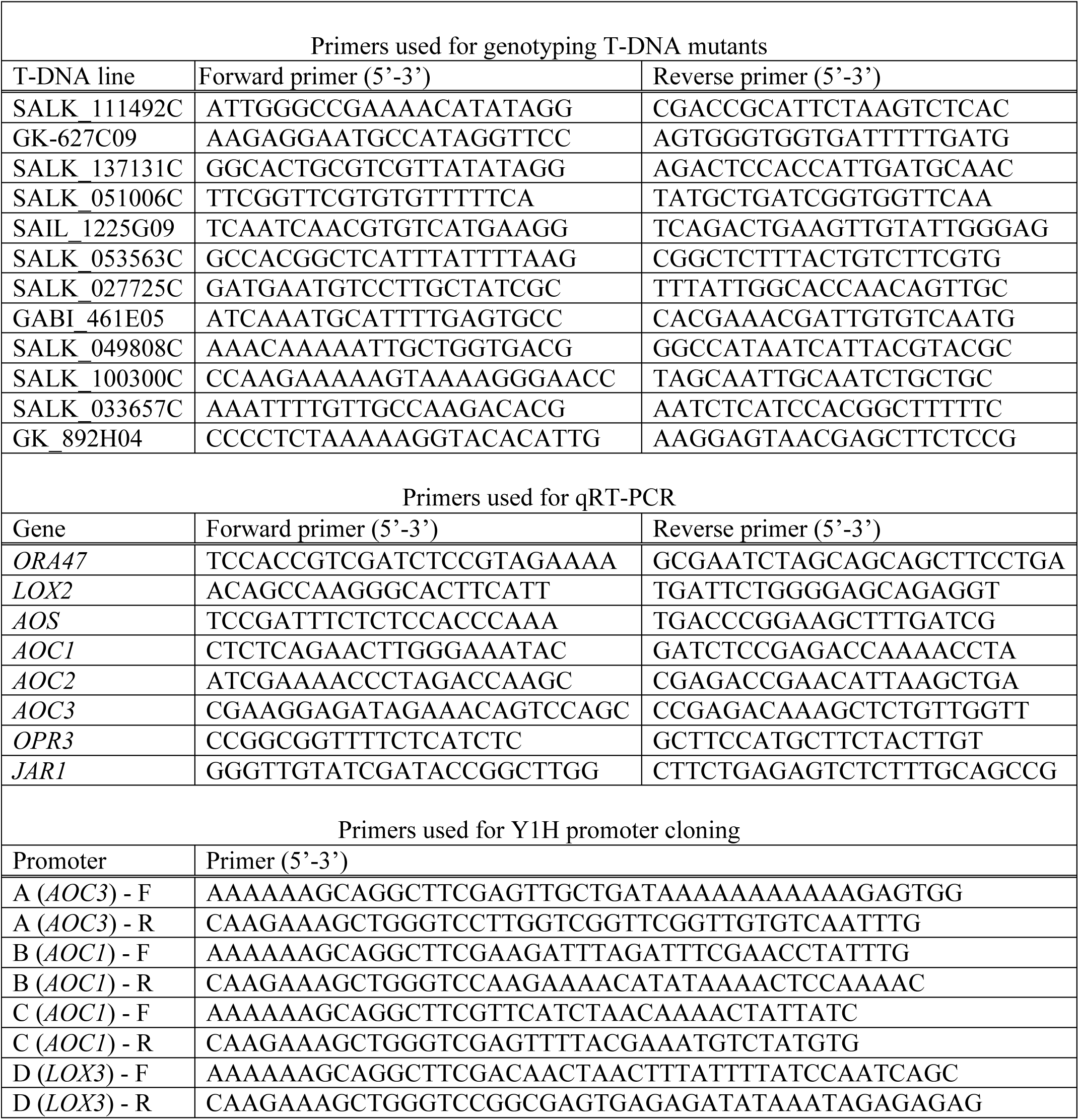
List of primers used for genotyping of T-DNA mutants, qRT-PCR analysis and promoter cloning for Y1H.

